# Isolating perceptual biases caused by trial history during auditory categorization

**DOI:** 10.1101/2022.01.17.476581

**Authors:** Daniel Duque, Jaime de la Rocha

**Author notes:** CORRESPONDING AUTHORS Daniel Duque; Jaime de la Rocha.

## Abstract

Just as most experiences have their origin in our perceptions, our perceptions can also be fundamentally shaped by our experiences. However, isolating which events in the recent past can impact perceptual judgments remains a difficult question, partly because post-perceptual processes can also introduce strong history dependencies. Two mechanisms have been hypothesized to specifically modulate perception: 1) the repulsive influence caused by previous stimuli and generally labeled as aftereffects, and 2) the modulation caused by stimulus predictions based on learned temporal regularities of the sensory environment, a key assumption in the predictive coding framework. Here, we ask whether these two mechanisms do indeed bias perception by training rats in an auditory task featuring serial correlations along the sequence of stimuli. We develop a detailed behavioral model that isolates the repulsive aftereffect generated by previous stimuli and shows that this repulsion cannot be explained from an interaction between past and current stimuli, and that it is still present in catch trials lacking the current stimulus. Moreover, the model describes that the bias caused by the animals’ expectation, as they leverage the predictability of the stimulus sequence, is present in a foraging task without the sensory component but with the same serial correlations in the sequence of rewards. These results indicate that the aftereffect and the prediction biases observed during an auditory task can all be revealed in the absence of a sensory stimulus, putting into question their perceptual nature.

## INTRODUCTION

The question of how previous decisions and outcomes affect future choices has been extensively investigated using dynamic foraging tasks (Sugrue et al., 2004; Samejima et al., 2005; Daw et al., 2006; Lau and Glimcher, 2008; Tai et al., 2012; Donahue et al., 2013; Kim et al., 2013; Groman et al., 2019; Moin Afshar et al., 2020). These are non-perceptual free-choice tasks where subjects have to choose between several options and where the probability of being rewarded after choosing each option varies with time. Despite the various versions of this type of task, in general subjects track the value of the different options using the history of previous choices and their outcomes and decide in each trial for one option based on these choice values. Thus, if presented with only two options (two-alternative forced choice task; 2AFC), subjects predominantly choose option A while being the most profitable, then they start opting for option B when the choice values cross (Zentall, 2020). The tracking of choice values may reflect the dynamics of ongoing Reinforcement Learning and may, for this reason, be an inevitable process at play during any type of reward-based decision-making task.

The neural bases of perceptual decisions, on the other hand, have been studied in psychophysics using 2AFC discrimination tasks in which the value of the two options is generaly constant and balanced (Newsome et al., 1989; Romo et al., 1998; Uchida and Mainen, 2003; Znamenskiy and Zador, 2013). Nevertheless, animals still develop sub-optimal trial history biases, the most common of which is a tendency to repeat previous choices, present even in the absence of trial feedback (Akaishi et al., 2014; Urai et al., 2017; Braun et al., 2018). In the presence of trial feedback, this repeating bias can depend on the trial outcome, usually becoming an attraction towards previous rewarded choices but a repulsion away from unrewarded ones (Fründ et al., 2014; Abrahamyan et al., 2016; Fan et al., 2018; Tsunada et al., 2019), a win-stay/lose-switch strategy similar to the one found in the foraging tasks. On top of these seemingly unavoidable previous choice/outcome biases, the magnitude of the previous stimuli, which is strongly correlated with previous choices, can also exert either repulsive (Addams, 1834; Holland and Lockhead, 1968; Cross, 1973; Bliss et al., 2017; Fritsche et al., 2017; Stein et al., 2020) or attractive biases (Fischer and Whitney, 2014; Papadimitriou et al., 2015; Bliss et al., 2017; Akrami et al., 2018; Barbosa et al., 2020; Stein et al., 2020). The attraction biases caused by previous stimuli are thought to be post-perceptual and supposedly arise from the maintenance of stimulus information in working memory (Bliss et al., 2017; Fritsche et al., 2017; Stein et al., 2020). Repulsive aftereffects on the other hand have been historically grounded in sensory adaptation (e.g., visual aftereffects: Thompson and Burr, 2009; auditory masking: Oxenham and Plack, 1998). According to this large body of work, the presentation of a particular stimulus causes neural adaptation or fatigue that transiently reduces the responsiveness to subsequent presentations of the same stimulus (Malone et al., 2002; Ulanovsky et al., 2003; Duque et al., 2016). This response reduction is what ultimately biases choices away from the choice associated with the adapted stimulus (Barlow and Hill, 1963; Kashino and Nishida, 1998; Dahmen et al., 2010). Despite the extensive evidence supporting this view, it is unclear if all repulsive biases are truly caused by sensory adaptation and in particular whether those observed in trial-based auditory 2AFCs fall into this category.

Another aspect that can modulate perception and bias perceptual judgments are the expectations about the incoming stimuli the brain can generate using an internal statistical model of the environment (Helmholtz, 1866; Rao and Ballard, 1999; Friston, 2005; Clark, 2013). In the framework of predictive processing, sensory circuits combine bottom-up stimulus inputs with top-down predictive signals elaborated in areas higher in the processing hierarchy (de Lange et al., 2018; Keller and Mrsic-Flogel, 2018). Expectations modulate the stimulus evoked responses along the sensory pathways (Kok et al., 2014; Carbajal and Malmierca, 2018) and bias perceptual judgments both in health and disease (Barrett and Simmons, 2015). Several studies have recently used a variation of a 2AFC task that, by including across-trial correlations in the sequence of stimuli, promotes the development of a predictive bias named *transition* bias: a tendency to repeat or alternate the previous response based on an internal estimate of the repeating probability of the sequence (Goldfarb et al., 2012; Jones et al., 2013; Meyniel et al., 2016; Kim et al., 2017; Braun et al., 2018; Hermoso-Mendizabal et al., 2020). Because the stimulus sequence is identical to the sequence of rewarded options, it is unclear whether the transition bias reflects a prediction about the stimulus or about the rewarded response. Based on human psychophysics, it has proposed that the transition bias is related to the processing of the stimulus and hence it should be classified as a perceptual bias (Wilder et al., 2009; Jones et al., 2013). We have recently shown that rats also display the Transition bias during an auditory 2AFC task with the same type of serial correlations (Hermoso-Mendizabal et al., 2020). However, direct evidence addressing whether it affects perception is still lacking.

Here, we investigate if the repulsive aftereffect and the predictive transition bias observed in an auditory 2AFC task are indeed affecting the perception of the current stimulus. We fit a behavioral model to the rats’ choices and show that there is no interaction between the previous and the current stimuli, indicating that the repulsive aftereffect does not have a perceptual impact. We confirm this by showing that this repulsive bias exists even in catch trials without sensory input. Furthermore, we show that rats develop a qualitatively similar predictive transition bias in both the Auditory task and in a Foraging task with no sensory stimuli, questioning its perceptual nature. Together, our results suggest that the repulsive aftereffect observed in our task may be caused by adaptation of post-perceptual circuits and that the transition bias reflects the prediction of the rewarded response rather than an expectation of future stimuli.

## RESULTS

### Previous stimuli caused a repulsive bias which did not interfere with the current stimulus

To characterize the impact of previous stimuli on the animals’ perceptual judgments, we investigated the behavior of rats (Group #1, n = 29) trained in an auditory two-alternative forced-choice task (2AFC) previously developed to explore expectation-based biases (Hermoso-Mendizabal et al., 2020). The task required rats to discriminate the interaural level difference (ILD), defined as the difference in intensity between two sounds presented in the Right and the Left speakers, to identify the side with the loudest sound, and seek reward in the associated port (Auditory task: Pardo-Vazquez et al., 2019; Hermoso-Mendizabal et al., 2020). To quantify the various factors influencing animals’ behavior, we fitted a generalized linear model to the sequence of choices of each rat (Net evidence GLM; Fig. 1A, see Methods). This GLM linearly combined (1) the samples of ILD from the *current stimulus*; (2) the ILD from each of the *previous stimuli* presented over the last ten trials, and (3) *previous rewarded* and *unrewarded responses*, i.e., positive and negative reinforcers (*r^+^* and *r^−^*). Including all these different history regressors aimed to isolate the impact of Previous stimuli from that of previous correct and incorrect responses, history events which are generally very correlated (see below).

**FIGURE 1.**
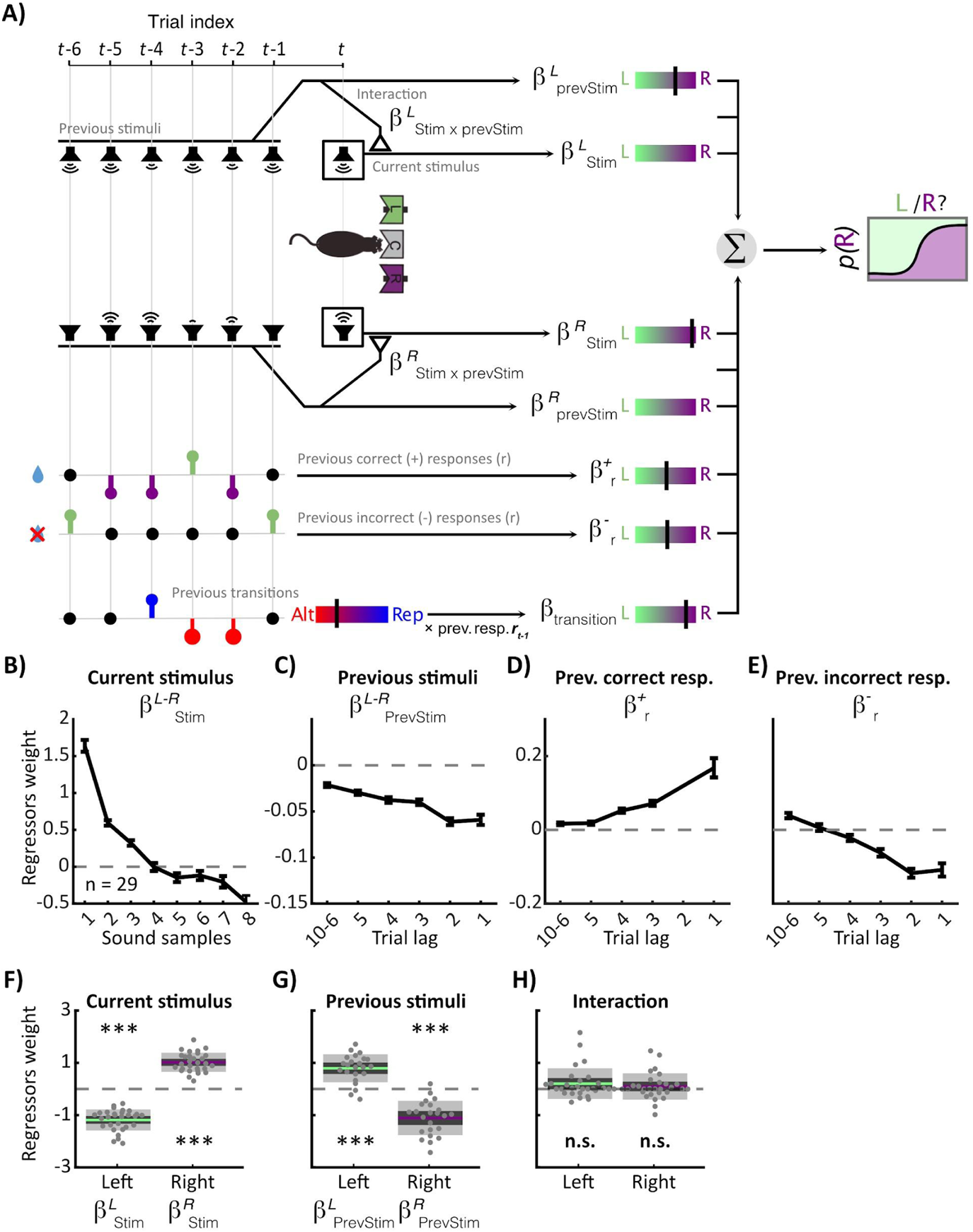
GLM analysis in a perceptual 2AFC Auditory task in rats. **A)** Exemple series of recent trials used to model rat responses in the current trial *t*. Rats’ responses were a function of the *Current stimulus* ILD, the ILDs from *Previous stimuli*, the side-specific interaction of previous stimuli with the current stimulus (*Interaction* term), previous *rewarded* (r+) and *non-rewarded* (r−) responses, and by previous *Transitions* (Repetitions, +1 and Alternations, −1). All these variables are weighted by different βs and linearly combined to generate the probability of a Rightwards response. **B-E)** Weights from the Net evidence GLM fitted without the interaction term for the Current stimulus (B; one point per stimulus sample), previous stimuli (C; one point per trial lag), and previous rewarded r^+^ and unrewarded r^−^ responses (D and E, respectively). Points show average weights for n = 29 rats (Group #1) and error bars show s.e.m. **F-H)** Weights from the Interaction GLM for the Current stimuli β^X^_Stim_ (F; ttest: all p < .0001, Cohen d > 2.5), Previous stimuli β^X^_prevStim_(G; all p < .0001, Cohen d > 1.5), and the interaction terms β^X^_Stim × prevStim_ (H; β^L^_Stim × prevStim_: p = .0714, Cohen d = 0.3; β^R^_Stim × prevStim_: p = .2946, Cohen d = 0.2).

The decisions of the animals were positively and strongly modulated by the Current stimulus (Fig. 1B; for a full report of the GLM see Fig. 1 - Supplement 1). Previous stimuli on the other hand, had a negative and long lasting impact on choices (previous stimulus *repulsive* bias; Fig. 1C). This dependence implies that, when in previous trials there was a strong sound intensity difference favoring one side, the upcoming choices were biased towards the opposite side. Importantly, this repulsive bias had the opposite effect of previous correct r^+^ and incorrect choices r^−^, which induced the common *win-stay-lose-switch* bias (Fig. 1D-E). Re-parametrizing previous responses r^+^ and r^−^ as Previous responses (*r^+^* − *r^−^*) and Previous reward side (*r^+^* + *r^−^*) illustrated more clearly that the repulsive effect of the magnitude of the Previous stimuli was the opposite to the stronger but attractive effect of Previous reward side (Fig. 1 - Supplement 2; see Methods). Teasing apart these opposite effects was possible due to the task design and the GLM parameterization: there were four mean ILDs within each Previous rewarded side, and each mean ILD stimulus was composed of a variable number of samples randomly drawn from a given distribution. This provided a large ILD variability within each Previous rewarded side for their impact to be separately assessed from that of the magnitude of the Previous stimuli. Moreover, we powered the fitting of the model using a total of 686,613 trials (an average of 27,119 trials per animal; range 6,433 - 55,792), allowing for a fine quantification of the impact of these different history events. Together, these opposite effects imply that e.g. after a Leftward correct choice there is a tendency in the next trials to choose Left, but that this attraction bias is weaker the stronger the previous stimulus is, as the attraction is partly compensated by the previous stimulus repulsion. This repulsive bias caused by the past presentation of sounds, independently of the response of the animal, is reminiscent of the aftereffect bias caused by sensory adaptation, a hypothesis we tested next.

**FIGURE 2.**
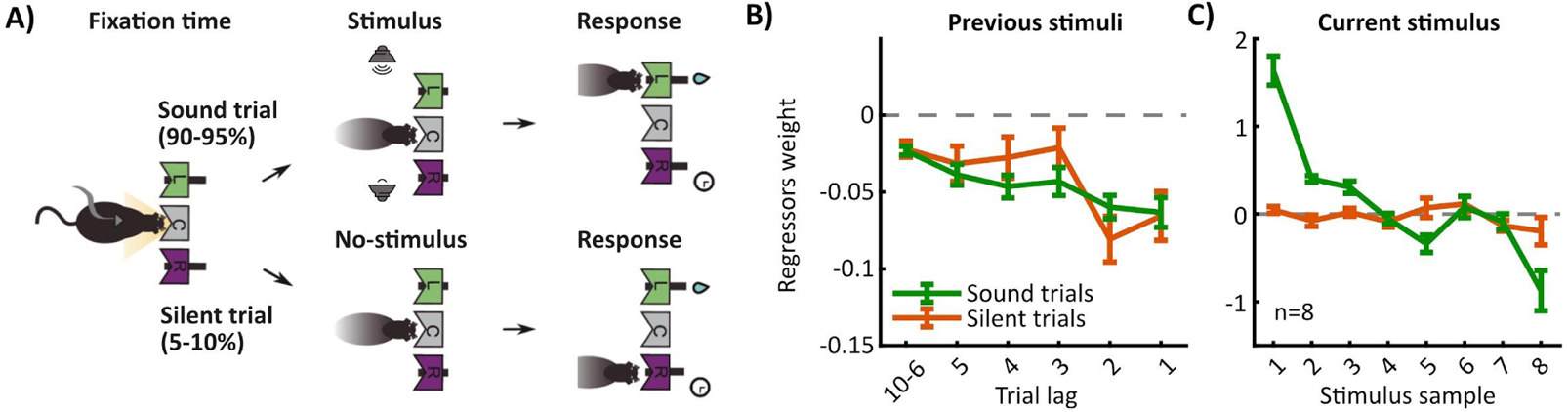
Previous stimuli cause a repulsive bias in Silent catch trials. **A)** Scheme of the 2AFC task with Silent catch trials. After fixating for 300 ms in the central port, in 90-95% of the trials, a stimulus is presented from two speakers and rats have to discriminate the side with the loudest sound. In the remaining 5-10% of trials, no stimulus is presented to guide the choice of the rat. **B)** GLM weights for Previous stimuli are not different between Sound and Silent trials (3-way ANOVA evaluating the weight of the Previous Stimuli kernel with ‘trial type’ as categorical-, ‘lag’ as continuous- and ‘animal’ as a random variable; ‘trial type’ main effect: F_1,85_ = 0.17, p = .6845; ‘trial type’ × ‘lag’ interaction: F_1,85_ = 0.65, p = .4215). **C)** Weights for the Current Stimulus. Points show average weights for n = 8 rats (Group #2) and error bars show s.e.m.

If the mechanism underlying the repulsion was for instance sensory adaptation, then the presentation of a loud sound on one side would decrease the sensory response of the following stimuli presented on that same side. This would in turn imply that the history of previous stimuli can cause an imbalance in the impact of the current Left and Right stimuli on the current choice (Fig. 1 - Supplement 3). To investigate this possibility, we extended our GLM to include an interaction between previous and current stimuli (Interaction GLM, Fig. 1A). To explicitly capture the interaction, we modified the GLM in three ways (see Methods): first, we simplified the Current stimulus regressor, previously defined for each sample (Fig. 1B), to a single regressor given by a weighted sum of the samples listened (Fig. 1E). The Previous stimuli regressor (Fig. 1C) was similarly recapitulated into a single regressor given by a weighted sum over previous trials (Fig. 1F). Second, we separated the contributions of the net evidence of the Current and Previous stimuli into two components each, one for the Right speaker (CurrentStim_R_ and PrevStim_R_) and one for the Left speaker (CurrentStim_L_ and PrevStim_L_). Third, we added interaction terms for each side that could capture the negative interaction predicted by the repulsive bias (CurrentStim_R_ × PrevStim_R_ and CurrentStim_L_ × PrevStim_L_; Fig. 1A). The fit of the Interaction GLM showed that, as before, Current and Previous stimuli contributed positively and negatively to the choice, respectively (Fig. 1E-F). However, we found that there was not a significant interaction between the Current and Previous stimuli (Fig. 1G). This result suggests that the repulsive bias observed in this task does not arise from an interference between previous stimuli and the current stimulus.

**FIGURE 3.**
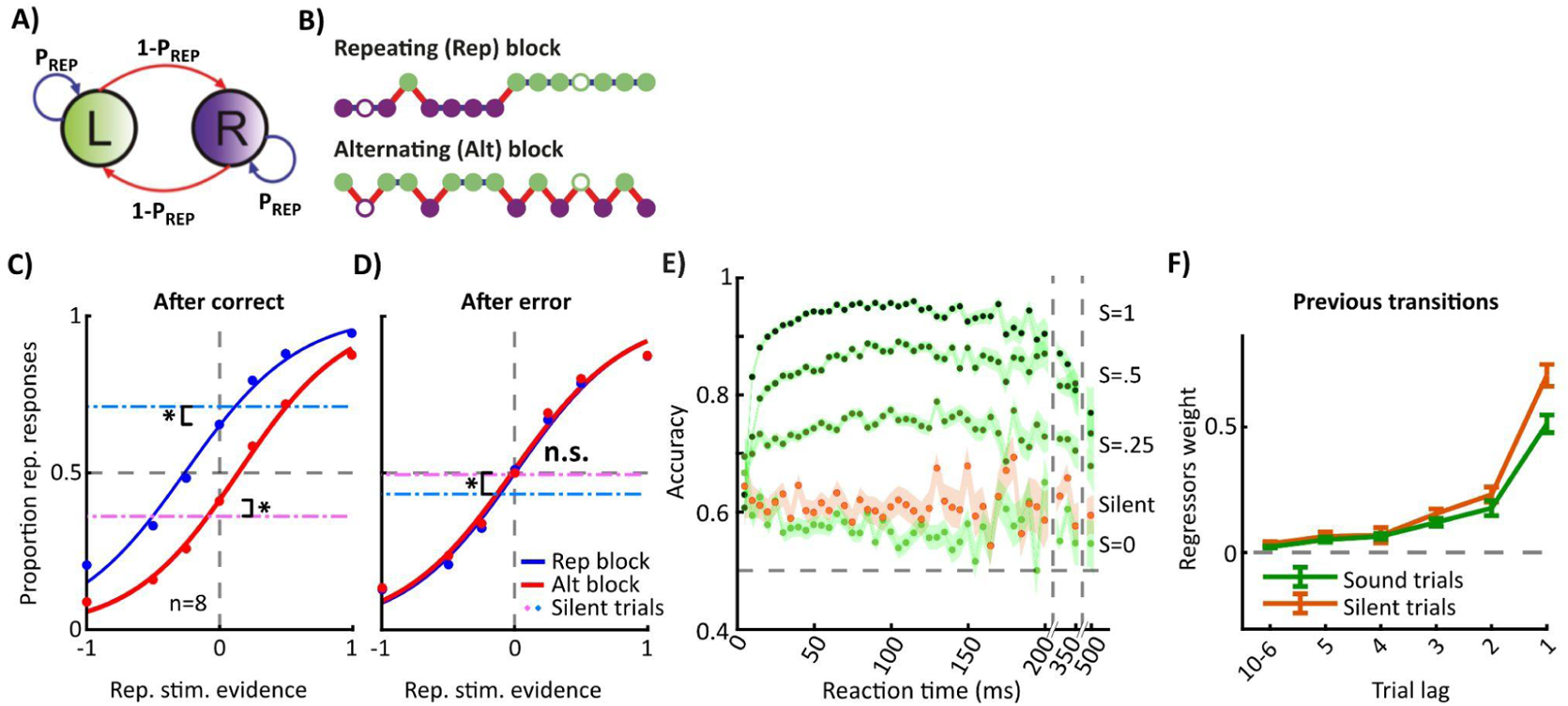
Repeating bias in Sound vs Silent catch trials. **A-B)** The sequence of stimulus categories Left/Right was generated using a two-state Markov chain parametrized with the probability to repeat the previous category P_REP_. which varied in blocks of trials generating repetitive (Rep) and alternating (Alt) blocks. Filled dots show Sound trials, while empty dots denote Silent trials. **C-D)** Mean psychometric curves showing the proportion of repeated responses vs. repeating stimulus evidence computed in trials following a correct **(C)** or an error response **(D)**, sorted by blocks (blue: Rep; red: Alt; n=8 rats in Group #2). After correct choices, the repeating probability in Silent trials (horizontal lines) was larger than *S*=0 Sound trials in Rep blocks (paired t-test, p = .0069) and lower in Alt blocks (p = .0072). After errors, the repeating probability was not different between Silent trials and *S*=0 trials in the Rep block (p = .4383), but it was in the Alt block (p = .0293). **E)** Tachometric curves showing the average accuracy versus reaction time (RT) conditioned on different stimulus strengths *S* (see *S-*values next to each curve) and for Silent trials (orange) for after-correct trials (RT bins were 5 ms below 200 ms; 50 ms between 200 and 350 ms; and 150 ms afterwards). Accuracy remained constant across RTs for Silent trials while it progressively decreased for zero evidence Sound trials (ANCOVA for ‘accuracy’ with ‘Stim. condition’ (*S*=0/Silent) as categorical- and ‘RT’ as a continuous variable; ‘Stimulus’ × ‘RT’ interaction F_1,35692_ = 6.08, p = .0137). **F)** Average transition weights were significantly larger for Silent trials than for Sound trials (3-way ANOVA for transition kernel with ‘Stim. condition’ as categorical-, ‘lag’ as continuous- and ‘animal’ as a random variable; main effect of ‘Stim. condition’: F_1,85_ = 6.79, p = .0108; ‘Stim. condition’ × ‘lag’ interaction: F_1,85_ = 3.75, p = .056).

### Stimulus repulsive bias is observed in Silent catch trials

Our previous model-based analysis suggests that the repulsive bias was not produced by a linear interference of previous stimuli on the perceptual impact of the current stimulus. But there could be more complex non-linear interactions between previous and current stimuli that our model did not capture. To have a more direct and conclusive test, we trained animals in Group #2 (n = 8 rats) in the same task but in a random 5-10% of catch trials, animals did not receive any acoustic sensory information (Fig. 2A, Silent trials). In the rest of the trials (Sound trials) stimulus evidence varied randomly from trial to trial as before (see Methods), and animals responded in Silent trials with a speed comparable to Sound trials (Hernández-Navarro et al., 2021). If the repulsive bias was truly not carried by the current stimulus, then it should be present in Silent trials. We fitted the Net evidence GLM separately for Silent and Sounds trials and found no significant difference in the Previous stimuli kernel (Fig. 2B), implying that the repulsive bias was equally present in both conditions. Furthermore, we confirmed that this was also the case when comparing Sound trials with accidental silent trials in a separate group of animals (Group #6, n = 18 rats; i.e. trials in which the reaction time was so short that the set-up did not have time to produce any sound: Fig. 2 - Supplement 1). Altogether, these results indicate that the repulsive bias caused by previous stimuli is instantiated independently of the value or the existence of the current stimulus, suggesting it is not the consequence of a perceptual aftereffect caused by sensory adaptation.

### Silent trials show more repeating bias than Sound trials in a 2AFC Auditory task

Having shown that the repulsive bias originates in previous stimuli but does not necessitate the current stimulus to impact choices, we next investigated the impact of expectations on the perception of the stimuli. To promote and control the use of subjects’ expectations, we had made the sequence of stimulus categories (right/left) partially predictable by introducing un-cued blocks of 80 trials with a tendency to repeat the previous stimulus category (Rep block; prob. to repeat P_REP_ = 0.8) or to alternate between the two stimulus categories (Alt block, P_REP_ = 0.2; Fig. 3A-B; Hermoso-Mendizabal *et al* 2020). As reported in a previous work (Hermoso-Mendizabal *et al* 2020), we found that animals capitalized on the predictability of the sequence and biased their perceptual choices accordingly: responses showed a tendency to repeat their previous choice in Rep blocks and to alternate it in Alt blocks (Fig. 3C). To investigate how this prediction bias affected the stimulus processing, we used again Silent catch trials in which the rewarded side followed the same Repetition or Alternation pattern but the associated sound was not presented. Accuracy was larger for Silent trials than for Sound trials with zero stimulus strength (ambiguous Sound trials; *S*=0): a 3-way ANOVA for ‘accuracy’ with variables ‘block’ (Rep/Alt), ‘Stim. condition’ (Sound/Silent) and ‘animal’ as a random factor yielded a significant main factor ‘block’ (F_1,28_ = 10.71, p = .0028) and ‘Stim. condition’ (F_1,28_ = 12.31, p = .0015), but no interaction (F_1,21_ = 0.02, p = .89). Next, we defined the *repeating probability* as the probability that animals repeated their previous choice and compared it in Silent trials versus *S*=0 Sound trials. We found that, after correct choices, the repeating probability in both blocks was more markedly different from 0.5 in Silent trials compared with Sound trials (0.72±0.08 vs. 0.66±0.10 in Rep block and 0.37±0.06 versus 0.41±0.06 in Alt block). After error choices, the repeating probability reset to 0.5 for both conditions and both blocks (Hermoso-Mendizabal et al., 2020), except for Silent trials in the Rep block which was slightly below (Fig. 3D).

To understand this difference in prediction bias between Silent and *S*=0 Sound trials we leveraged on the design of the task in which the stimulus duration was determined in each trial by the reaction time (RT) of the animal (see Methods). We examined how accuracy changed with RT by plotting the tachometric curve (Fig. 3E; Stanford et al., 2010). Because of the repeating bias, accuracy started at the same above-chance level at RT=0 for both types of trial. However, accuracy remained constant as a function of RT for Silent trials as there was no sensory input to be integrated (Fig. 3E; orange line). In Sound trials, in contrast, it increased with RT for all stimulus strengths except for *S*=0, for which the accuracy decreased (Fig. 3E; green lines). This suggests that the combination of evidence coming from previous trials and from the stimulus occurred serially: the repeating bias set the baseline of a putative decision variable at stimulus onset, resulting in an above-chance accuracy at near zero RTs. As RT increases, stimulus evidence is integrated for *S*>0 yielding a higher accuracy. For *S*=0 Sound trials, because stimulus fluctuations are dissociated from the reward side, the integration of the stimulus mimics accumulation of noise, thus reducing the accuracy as RT increases.

To quantify the impact of the expectation built from the history of previous trials on choice we included regressors in the Net evidence GLM representing the previous correct Transitions, defined as pairs of consecutive rewarded responses in which the animal either made a Repetition (+1) or an Alternation (−1) between the two possible choices (Fig. 1A). Including previous Transitions in the model can capture the animals’ tendency to repeat or alternate the previous response leveraging the repeating and alternating statistics introduced in each block. We fitted the Net Evidence GLM separately in Sound and Silent trials following correct choices and found that the Transition kernel was significantly larger for Silent trials than for Sound trials (Fig. 3F). This difference is consistent with transitions explaining a lower fraction of the behavioral variance in Sound trials. Moreover, the after-error reset of the Transition kernel occurred in both Sound and Silent trials (Fig. 3 - Supplement 1: after-error). We finally found that the *building* of the expectation bias did not depend on the presence of the stimulus in past trials meaning that previous transitions containing Silent trials provided as much evidence to generate a certain expectation as the transitions containing only Sound trials (Fig. 3 - Supplement 2). In total, these results show that the accumulation of previous transitions, the after-error reset as well as the impact on choice of the transition evidence, are all mechanisms which occur independently of the presence of sensory inputs. In other words, animals seemed to use previously rewarded transitions, independently of the sensory evidence guiding those transitions, to predict future rewarded transitions but not to predict the upcoming stimuli.

### Animals develop a expectation bias in a 2AFC Foraging task

If the expectation bias can be both effective in Silent trials and built up from previous transitions containing Silent trials, could it be observed in the complete absence of stimuli? To answer this, we trained animals in Group #3 (n = 7 rats) in a novel 2AFC Foraging task which contained no sensory information to guide their choices (Foraging task). In this task, animals self-initiated each trial by poking in a central port, waited for 300 ms (fixation time) and then they guessed if the reward was in the left or the right port (Fig. 4A). Critically, as in our previous tasks, the sequence of rewarded sides was structured in repetitive and alternating blocks (Fig. 4B-C). Rats’ accuracy in both blocks was significantly above chance (Fig. 4D; mean±SD of 0.63±0.02 in the Rep block; 0.58±0.05 in the Alt block) indicating that animals were able to leverage the serial correlations and adapt their history bias in each block to improve their task performance. After correct choices, the repeating probability was significantly larger than 0.5 in Rep blocks (0.83±0.07; Fig. 4E), while it was significantly smaller than 0.5 in Alt blocks (0.31±0.15; Fig. 4F). After-error trials however, it was not different from 0.5 (Rep: 0.48±0.07; Alt: 0.55±0.08) showing that the reset of the repeating probability persisted in this non-perceptual Foraging task (Fig. 4E-F). This implies that the reset strategy cannot be interpreted as if the rats were ignoring the transition evidence accumulated across trials to focus on listening to the sound after an error choice, but instead it suggests the existence of a more general mechanism independent of the presence of a stimulus (Molano-Mazón et al., 2021). The repeating probability was mainly driven by previous Transitions (Fig. 4G) while the contribution of the previous correct (r^+^) and incorrect responses (r^−^) was much smaller (Fig. 4H-I). Thus, animals in the Foraging task were able to develop the same predictive transition bias as in the Auditory task, but just based on previous rewarded transitions, demonstrating that this type of expectation bias is independent of the processing of the stimulus.

**FIGURE 4.**
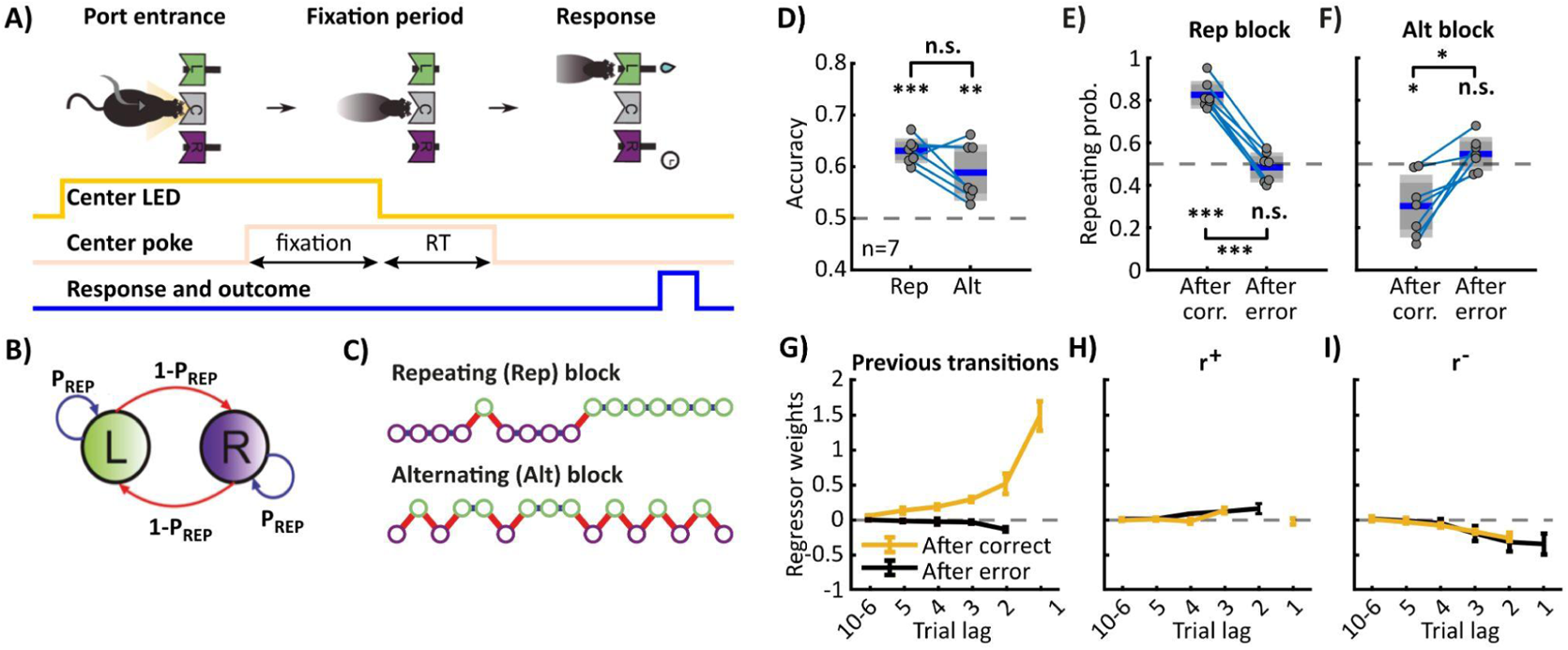
Transition bias in a 2AFC Foraging task. **A)** Scheme of the 2AFC Foraging task. Details of the task are identical to the Auditory task. At the end of the fixation period, the LED in the central port turned off indicating animals could seek reward in either the left or right port. **B)** The probability of repeating the previous stimulus was adjusted, **C)** generating Rep and Alt blocks. Empty dots denote Silent trials. **D)** Mean proportion of correct responses (n = 7) for Rep and Alt blocks. The accuracy is above chance (50%) in both blocks (Rep block, paired t-test p < .0001; Alt block, p = .0052; Rep vs. Alt was p = .09). **E-F)** Repeating probability in Rep **(E)** or Alt blocks **(F)**, and sorted by performance in the previous trial (after correct/after error). The repeating probability in a Rep block is larger than 0.5 (t-test, p < .0001), while in the Alt block is smaller than 0.5 (p = .0135), demonstrating rats can extract both sequential correlations. The repeating probability in after-error trials was not different from 0.5 (Rep: p = .55; Alt: p = .18). In both blocks, the repeating probability was different in after-correct versus after-error trials (paired t-test; Rep: p < .0001, Alt: p = .0059). **G)** Influence of past events on current choice after a correct (yellow) or error trial (black). Details as in Figure 3. Predictive transition bias guides the behavior of the animals. After-error choices, transition bias goes to zero. The impact of previous correct (r^+^) and incorrect responses (r^−^) is much smaller.

### Transition bias is larger in the Foraging task than in Silent catch trials

To make a quantitative comparison between the two tasks we next compared Silent catch trials in the Auditory task with the trials in the Foraging task (Fig. 5; see also Fig. 5 - Supplement 1). To compare the response repeating probabilities across blocks and tasks, we defined the *Excess of Repeating Probability* (ERP, see Methods) as the difference between 0.5 and either the repeating probability (in the Rep block) or the alternating probability (in the Alt block). While after-correct trials the ERP was significantly larger for the Foraging task than for Silent catch trials (Fig. 5B, left) there was no difference in after error trials (Fig. 5B, right). This difference was caused because rats gave a larger weight to previous transitions in the Foraging task than in Silent trials (Fig. 5D). No differences were observed between tasks in the previous correct *r^+^* and incorrect *r^−^* response kernels (Fig. 5E). Altogether, this data suggests that in absence of sensory information, rats modulate the probability to repeat their previous choice by means of the predictive transition bias. Intriguingly, such an increase in Transition bias is not observed when we modified other variables, such as the magnitude of the sequential correlations (Fig. 5 - Supplement 3).

**FIGURE 5.**
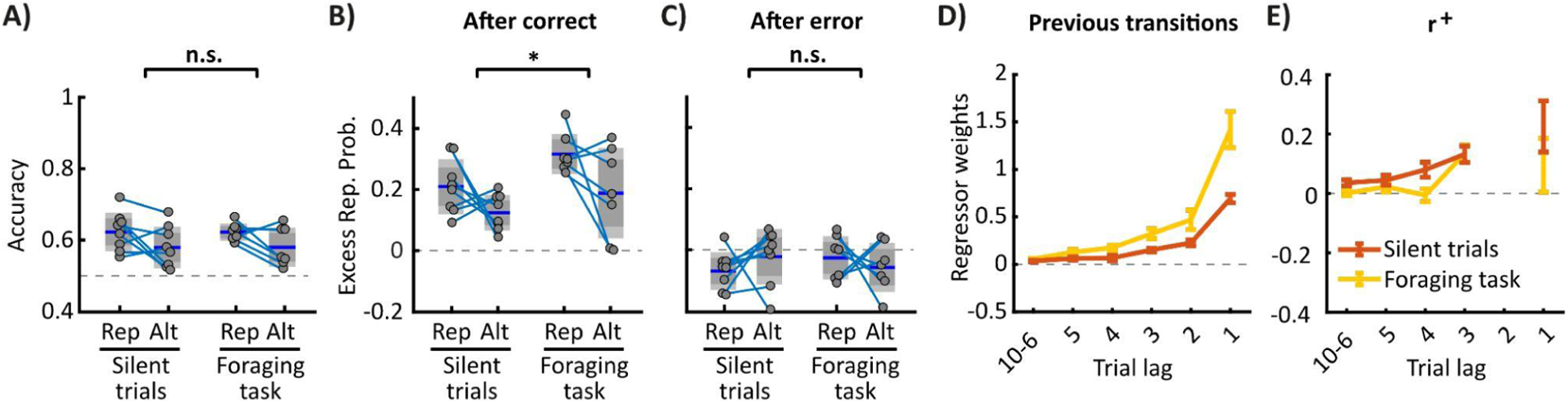
Repeating bias comparison between the Foraging and Silent catch trials. **A)** Accuracy for Silent trials in the Auditory task and all trials in the Foraging task sorted by Rep (P_REP_=0.8) and Alt blocks (P_REP_=0.2) showed no significant difference (2-way ANOVA for ‘accuracy’ with ‘block’ and ‘task’ as variables; ‘block’ × ‘task’ interaction F_1,22_ = 0.57, p = .4585; ‘task’ F_1,22_ = 0.02, p = .8986). The accuracy for Rep and Alt blocks was also similar (‘block’ F_1,22_ = 3.28, p = .0838). **B-C)** Excess of repeating probability (ERP; i.e. repeating probability normalized to positive values) in after-correct **(B)** or after-error trials **(C)** sorted by type of trial (Silent/Foraging) and block (Rep/Alt). In after-correct trials (B), the ERP is significantly larger in the Foraging task than for Silent trials, and in Rep compared with Alt blocks (2-way ANOVA; ‘task’ F_1,22_ = 8.18, p = .0091; ‘block’ F_1,22_ = 4.36, p = .0486; ‘block’ × ‘task’ interaction F_1,22_ = 0.64, p = .4306). **C)** No difference in ERP was observed after-error trials (‘task’ F_1,22_ = 1.64, p = .2142; ‘block’ F_1,22_ = 1.07, p = .3126; ‘block’ × ‘task’ interaction F_1,22_ = 0.29, p = .5928). **D-E)** Average GLM transition kernels for Silent trials (orange) were smaller than for the Foraging task (yellow; ‘task’ F_1,92_ = 21.73, p < .0001; ‘task’ × ‘lag’ interaction F_1,92_ = 11.26, p = .0012). **E)** Weights for previous correct responses show no difference between Foraging task and Silent trials (‘task’ × ‘lag’ interaction F_1,76_ = 0.32, p = .571; ‘task’ F_1,76_ = 1.47, p = .2292). Weights for previous incorrect responses also show no difference (not shown: ‘task’ × ‘lag’ interaction F_1,76_ = 0.52, p = .4729; ‘task’ F_1,76_ = 1.79, p = .1851).

### Stimuli can accelerate the updating dynamics of the transition bias

Although animals implemented the predictive transition bias based on previous transitions independently of the sensory information, the stimuli had a great impact on accelerating the trial-by-trial updating of the transition evidence. To show this, we compared the transient dynamics of the accuracy and repeating probability during a block switch in the Auditory and the Foraging tasks (Fig. 6). In the Auditory task, baseline accuracy was relatively high and the stimuli helped animals maintain their accuracy almost unaffected during the block switch. For that reason, animals could experience a high rate of correct transitions, i.e. transitions made of two consecutive reward choices, which allowed them to update their repeating bias rapidly (Fig. 6C). In contrast, in the Foraging task, baseline accuracy was lower and it dropped markedly and abruptly in the first trial of the block as animals had no way to guess the block switch. Accuracy then remained low for many trials, dragging the updating of the repeating probability which in turn slowed the recovery of the accuracy. As a result, both quantities took more trials to reach the block’s baseline in the Foraging task compared with the Auditory task (Fig. 6B,D). Altogether, this reflects that, even though animals developed the same transition bias independently of the task, the presence or absence of stimuli can catalyze or slow the accumulation of repeating evidence which, because it is based only on correct transitions, is very dependent on the overall animals’ accuracy.

**FIGURE 6.**
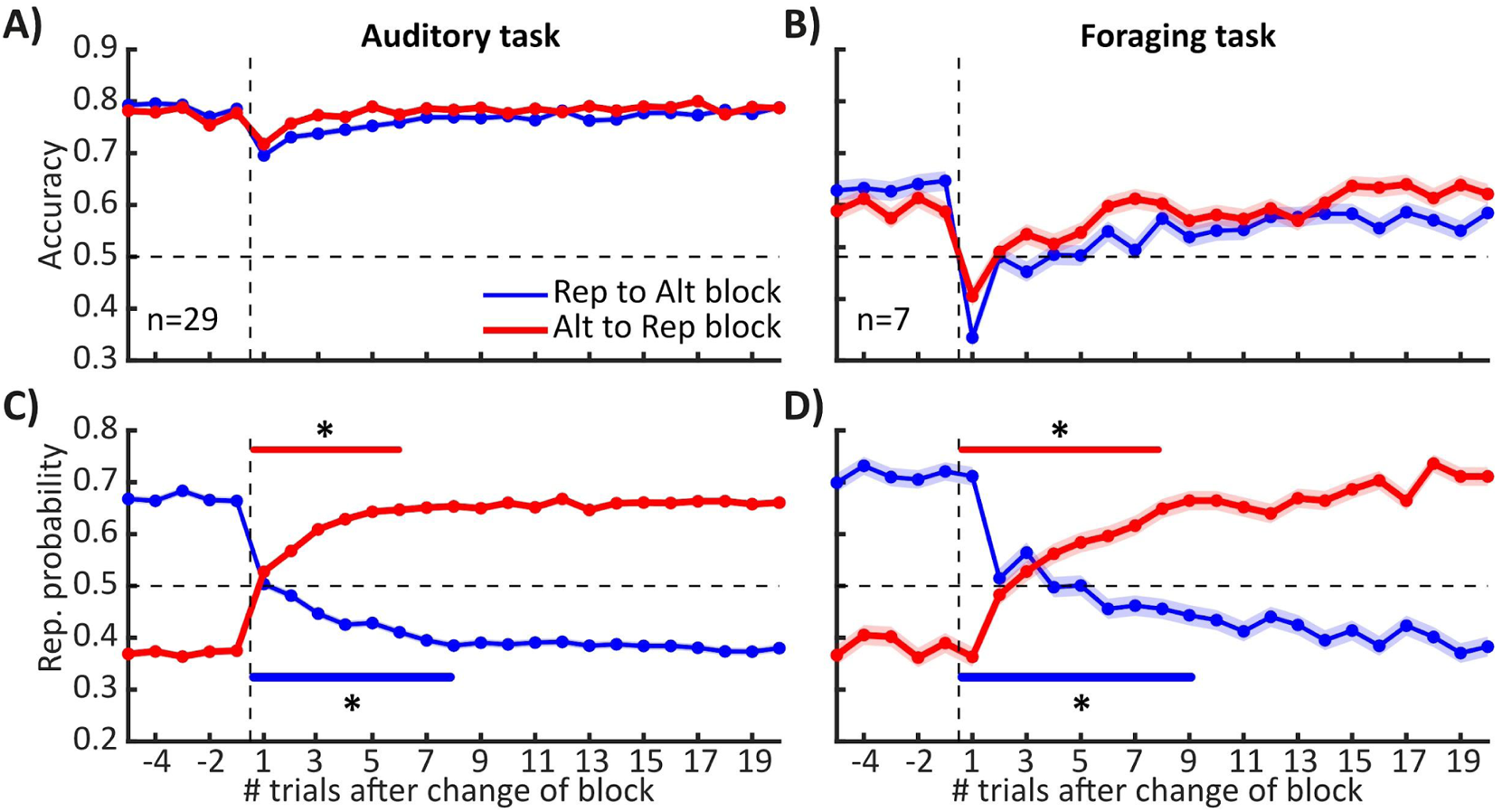
Updating dynamics at the block changepoint in the Auditory and Foraging tasks. **A-B)** Average accuracy in a change of block from a Rep to an Alt block (blue) and from an Alt to a Rep block (red) in the Auditory task (A) and the Foraging task (B). **C-D)** Repeating probability in a block change in the Auditory (C) and the Foraging task (D). The number of trials needed to recover the baseline repeating probability was greater in the Foraging task compared with the Auditory task. Horizontal lines in C-D show trials with a repeating probability different from baseline (ttest, p>0.05).

## DISCUSSION

In this study we investigated whether perception can be impacted by previous stimuli or by stimulus expectations built from the temporal regularities of the stimulus sequence. First, we found that the repulsive bias caused by previous stimuli does not modulate the impact of the subsequent stimuli, and it is present even in catch trials with no stimulus. Second, we found that animals performing a non-perceptual 2AFC foraging task with the same sequential correlations still develop the predictive transition bias, suggesting that this type of bias is predicting the rewarded transitions rather than the stimuli associated with it.

### Repulsive aftereffects during a perceptual 2AFC task

Repulsive sensory aftereffects were first described in the visual system almost two hundred years ago in the well known motion aftereffect (Addams, 1834). The most prevalent explanation for such aftereffect bias has been sensory adaptation (Barlow and Hill, 1963): repeated stimulation of sensory neurons with a specific stimulus input reduces the neuronal response to subsequent similar inputs. In the auditory system, repulsive stimulus aftereffects are observed for a plethora of acoustic properties such as intensity (Holland and Lockhead, 1968; Cross, 1973), direction of frequency modulation (Kay and Matthews, 1972), sound localization (Kashino and Nishida, 1998), azimuth motion (Grantham, 1998) or the ventriloquism aftereffect (Recanzone, 1998). In the intensity domain in particular, the reduction of perceived loudness is maximum when the preceding sound is louder than the subsequent sounds (Scharf et al., 1983; Marks, 1994), while when the sounds have similar intensities the reduction is smaller (Mapes-Riordan and Yost, 1999). Putative neuronal correlates of the aftereffect bias have been identified in the auditory system, and are also linked to sensory adaptation (Ulanovsky et al., 2003; Duque et al., 2016). Recent work has shown that sensory adaptation has indeed a perceptual impact on animals performing a frequency discrimination 2AFC task (Gronskaya and von der Behrens, 2019). However, in this study the repulsive effect was caused by an “adaptor stimulus” presented in the same trial right before the stimulus, as opposed to the across-trial sequential analysis that we performed. Addressing after-effects caused by previous stimuli in the sequence is challenging because one must isolate their effect from that of previous responses and outcomes. Sequential effects have been previously described in humans performing subjective reports of perceived acoustic intensity (for a review see DeCarlo and Cross, 1990), where a common pattern found was attraction to the previous response and repulsion to the previous stimulus. Here, we report a similar effect in rats performing a categorical 2AFC task except that we circumscribe the attraction to only correct responses and statistically resolve the time scale of the effects many trials back. In these previous reports, the repulsive intensity aftereffect was assumed to be caused by a negative interaction of previous stimuli on the perception of future sounds (Cross, 1973; Ward, 1973). However, none of these studies tested for the existence of that interaction. Our results show for the first time that the repulsion caused by previous stimulus does not interact with the impact of current stimuli (Fig. 1H) as well as it is also present in Silent catch trials (Fig. 2B), suggesting that this effect is not produced by loud sounds masking the perception of forthcoming sounds.

What kind of mechanism could explain this repulsive bias which, caused by previous stimuli, can be manifested in the absence of sensory input? (Fig. 7A). The most parsimonious mechanism underlying any repulsive bias is the response adaptation of neurons whose firing is modulated by the intensity of each of the two sounds. Neuronal adaptation, an ubiquitous feature along the auditory pathway (auditory nerve: Nomoto et al., 1964; inferior colliculi: Malmierca et al., 2009; auditory thalamus: Anderson et al., 2009; auditory cortex: Ulanovsky et al., 2003), is typically described as a decrease in the gain of the rate-level function, defined as the stimulus evoked firing rate versus stimulus intensity (Fig. 7B). This divisive adaptation would however predict an interaction between previous and current stimuli that we did not observe, as the impact of the current stimulus is determined by the gain of the rate-level function which is affected by previous stimuli (Fig. 7B). Moreover, the firing of these sensory neurons in Silent catch trials would be independent of the level of adaptation and hence could not explain the presence of the repulsion we found in these trials (Fig. 2D). An alternative possibility would be that adaptation is not divisive but subtractive. Subtractive adaptation implies a constant decrease in responsiveness along the rate-level function, and hence requires a sufficient baseline firing rate for the effect to be purely subtractive (Fig. 7C). If auditory sensory neurons were adapted by previous stimuli in an subtractive way, this would produce no interaction between previous and current stimuli, because the gain of the rate-level functions would remain the same, but would require baseline firing rates to be modulated by previous sounds to explain the repulsion found in Silent catch trials (Fig. 2D). To the best of our knowledge, pure subtractive adaptation has not been observed in auditory neurons, posing into question this mechanistic explanation. A final possibility would be that adaptation takes place in decision making circuits up in the hierarchy from sensory neurons (Fig. 7E). Current models of decision making postulate that circuits in parietal and frontal cortices receive inputs from sensory areas and, operating in a winner-take-all regime, establish a competition between Left- and Right-choice neurons which ultimately determines the current decision (Wang, 2002; Fusi et al., 2007; Roxin and Ledberg, 2008; Wimmer et al., 2015; Prat-Ortega et al., 2021). The classic ramping-to-bound spiking activity observed in these areas during decision formation (Kim and Shadlen, 1999; Roitman and Shadlen, 2002; Thura and Cisek, 2014; Hanks et al., 2015) could in principle activate long-lasting adaptive currents in “winning neurons” which could unbalance the competition in future decisions (Fig. 7E). For this to be plausible three conditions should hold: first, the stimulus intensity should modulate the total number of spikes fired by these neurons above and beyond the obvious dependence caused by the generated choice, i.e. beyond the categorical modulation caused by the ramping up versus ramping down. In other words, the ramping of e.g. Left-decision neurons in Left-choice trials should exhibit a dependence on the stimulus intensity. Using behavioral modeling, we have recently shown that in this task, there is evidence accumulation but that reaction times are mostly determined by an internal stimulus-independent timing signal (Hernández-Navarro et al., 2021). This implies that in Left-choice trials, Left-decision neurons may show larger slopes when the Left stimulus is strong without change in the duration of the integration (i.e. reaction time). This would cause larger firing the stronger the stimulus within any given choice. Second, an independent mechanism should reinforce rewarded decisions and weaken unrewarded ones in order to generate the win-stay-lose-switch behavior (see e.g. (Fusi et al., 2007). Third, the neural circuits that implement choice selection should be the same in Sound and Silent trials. That way, the adaptation caused by previous stimuli can leave a trace that unbalances the competition in trials with no stimuli. Although we have no evidence of that the ramping activity observed in these circuits during stimulus categorization persists in Silent trials, there is indirect evidence showing that most of the ramping observed is not caused by the integration of the stimulus but by internal urgency signals which could be at play in both Sound and Silent trials (Churchland et al., 2008; Thura et al., 2012; Park et al., 2014). In that scenario, the driving input into the decision circuit would be coming from a non-sensory action initiation circuit (Fig. 7E) as postulated by behavioral modeling (Hernández-Navarro et al., 2021). If adaptation of decision neurons underlies the aftereffect, it would be a phenomenon with a sensory origin causing a non-perceptual decisional effect (Bosch et al., 2020). Electrophysiological experiments assessing whether the previous stimuli aftereffect can be observed in the activity of auditory sensory neurons or in decision-related brain areas will help elucidate whether any of the proposed mechanisms is at play (Macke and Nienborg, 2019).

**FIGURE 7.**
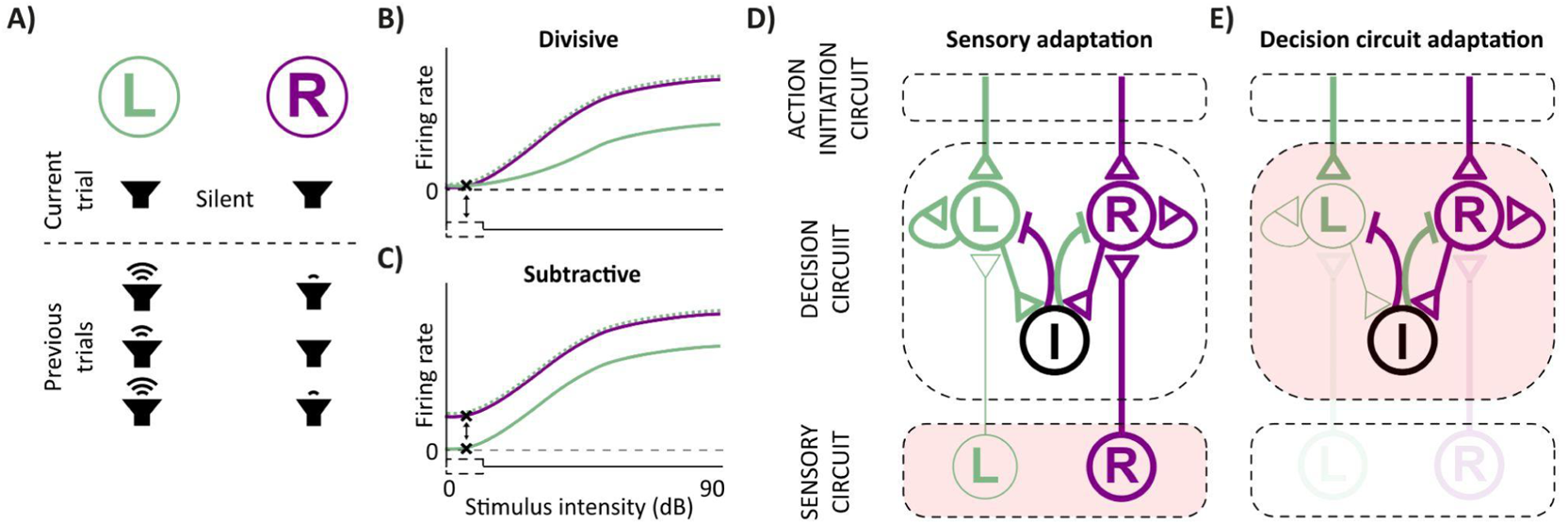
Theoretical models for the mechanism underlying the repulsive bias observed during Silent catch trials. **A)** Schematic of a Silent catch trial in which there is a history of previous Left loud sounds that creates a Left repulsion bias (i.e. a bias towards the Right), by reducing either the Left sensory responses (B-D) or the Left-choice population responsiveness at the Decision network (E). **B-C)** Rate-level functions showing response firing rate of Left and Right sensory neurons as a function of sound intensity. Because of previous Left loud sounds (A), Left-sensory neurons show a reduced response (compare adapted in solid with unadaptated as dotted). Such adaptation may be Subtractive **(B)** or Divisive **(C).** However, only subtractive adaptation would generate the Aftereffect bias in Silent trials (white box at 0 dB). **D-E)** Schematic of a network consisting of a sensory circuit with stimulus selective populations sending topographically organized inputs to a decision circuit with choice-selective populations. An action initiation circuit promotes choice formation by setting the competition between choice-selective populations via sensory-independent inputs. If the Sensory circuit shows subtractive adaptation, the Left input to the Decision circuit in a Silent trial would still be weaker compared to the Right input biasing the choice towards the Right (D). If the Decision circuit shows adaptation, in a Silent trial the sensory input will not play any role, and it would be a competition set by the action initiation circuit that would result in a Right biased decision (E).

### Silent catch trials in a perceptual 2AFC task

Early psychophysics studies included Silent catch trials in order to measure the influence of choice biases, but such practice was lost with time (White and Wixted, 2010). Here, we have shown that the accuracy of rats’ responses is better for Silent catch trials than for ambiguous *S*=0 Sound trials. Interestingly, the accuracy for *S*=0 Sound trials inversely correlated with reaction time; so the longer the reaction time the lower the accuracy. This was not the case for Silent catch trials, where the accuracy was constant irrespective of the reaction time (Fig. 3E). Experiments in monkeys and rats using a motion discrimination 2AFC task have shown that accuracy does not decrease with time for *S*=0 trials (Gold and Shadlen, 2003; Reinagel, 2013; Shevinsky and Reinagel, 2019). That difference can be explained by the fact that, as those classical tasks do not present serial correlations, the baseline accuracy at short reaction times is already at chance level. For the same reason, a 2AFC odor discrimination task also showed accuracy does not decrease with time for *S*=0 trials (Rinberg et al., 2006). However, as we observed for sound sampling, beyond a relatively short minimum, longer odor sampling times tended to decrease accuracy at increased reaction times (Uchida and Mainen, 2003). As such decrease does not occur for Silent catch trials, our data implies that the combination of the predictive transition bias with the sensory information occurred in a serial order: the transition bias sets a predetermined decision, then the stimulus is integrated guiding the final decision. In a drift diffusion model, that would represent that the transition bias sets the initial offset of the decision variable resulting in an above-chance accuracy at near zero RTs for both Sound and Silent catch trials. As RT increases, stimulus evidence is integrated for *S*>0 yielding a higher accuracy. For *S*=0 Sound trials however, the integration of the stimulus mimics accumulation of noise and hence reduces the accuracy as stimulus duration increases. For Silent catch trials, as no sensory input is ever integrated, the decision variable remains approximately constant (no sign of integration leak) and the accuracy does not decrease with RT. In the mechanistic model outlined above, the transition bias could be implemented by a transient prediction signal that is input into the decision circuit before the stimulus and is independent of the action initiation sustained urgency signal (Fig. 7).

### Animals do not predict stimuli but reward

In the framework of predictive processing, sensory circuits combine bottom-up stimulus information with top-down predictive signals (de Lange et al., 2018; Keller and Mrsic-Flogel, 2018). In a perceptual 2AFC task it is often assumed that the system predicts the forthcoming stimuli. However, the brain could be predicting 1) the stimulus category (louder on Left or Right speaker), which will evoke a *sensory* prediction error signal (Näätänen et al., 1978; Ulanovsky et al., 2003; Parras et al., 2017); 2) the reward side (Right or Left), which will evoke a *reward* prediction error signal (Daw et al., 2006; Tai et al., 2012; Lak et al., 2020a, 2020b); or 3) more complex reward patterns such as repetitions or alternations, which will evoke a *state* prediction error signal (Gläscher et al., 2010, Meyniel et al., 2016; Hermoso-Mendizabal et al., 2020). Thus, if the absence of the guiding sensory stimuli does not alter the impact of the prediction on choice, where does this prediction signal come from?

Predictive history choice biases can be first-order (e.g., ‘last trial I went to the *left’*), second-order (e.g., ‘last time I *repeated* my action*’*) or higher-order (e.g., ‘there is a hidden sequence *left-left-right* which is particularly rewarding’). First-order sequential biases, described extensively in both non-perceptual foraging tasks (Sugrue et al., 2003; Samejima et al., 2005; Daw et al., 2006; Lau and Glimcher 2008; Tai et al., 2012; Groman et al., 2019; Moin Afshar et al., 2020) and in 2AFC perceptual discrimination tasks (Busse et al., 2011; Abrahamyan et al., 2016; Urai et al., 2017; Fan et al., 2018; Tsunada et al., 2019), reflect the estimation of the base rate of either stimulus or reward values (Wilder et al., 2009; Meyniel et al., 2016). In second-order effects, the occurrence probability of the stimulus or reward values are irrelevant and what biases behavior is the estimate of the transition probabilities, e.g. the probability that after a Left stimulus there is another Left stimulus (Cho et al., 2002; Yu and Cohen, 2008; Goldfarb et al., 2012; Meyniel et al., 2016; Hermoso-Mendizabal et al., 2020). Previous reports suggest that rats can develop third-order history choice biases in a non-perceptual foraging task (Tervo et al., 2014), but their characterisation is far less studied than first and second order biases.

Previous research has proposed that first-order history choice biases were associated with the response processing and affect the execution of the action, while second-order history choice biases arise from the processing of previous stimuli and ultimately affect the perception of the current stimulus (Maloney et al., 2005; Wilder et al., 2009). This classification is based on the analysis of the onset latency of the lateralized readiness potential (LPR), a hemispheric asymmetry of activation in the motor cortex, that can be obtained by alignment to the stimulus or to the response onset thus reflecting each of the two processing stages (Coles, 1989). Wilder and colleagues (2009) showed that the response and the stimulus LPR latencies varied with the trial history consistently with first and second-order biases, respectively. Moreover, when reanalysing data from a task which removes the response in some trials (Maloney et al., 2005), they observed that the first-order sequential effects were essentially absent. Finally, Jones and colleagues (2013) also re-analyzed a previous experiment in which the stimulus was eliminated in some trials and found that reaction times variability could be described by first-order but not by second-order effects (Wilder et al., 2013). At odds with this proposal, we have shown that second-order history choice biases appear even in the absence of sensory cues, suggesting it’s main effect is not altering the perception of the current stimulus. The differences in methodology and species makes a direct comparison between experiments difficult. Future electrophysiological recordings in the auditory cortex of animals performing the task will help elucidate the extent to which auditory responses are modulated by the transition bias as well as whether activity in Silent trials show traces of this internal prediction.

### Reset after error does not prioritize integration of sensory information

Finally, we observed that animals performing a non-perceptual foraging task still present a reset of the Transition bias after error trials. The reset after error, i.e. the ability in which animals temporarily ignore the recent transition history and respond randomly after an error choice, was described in an auditory categorization task (Hermoso-Mendizabal et al., 2020). Other experiments have shown similar results where, following an error choice, subjects selected their next move less influenced by the context (Braun et al., 2018; Kikumoto and Mayr, 2019), although they did not dissociate between transition and win-stay/lose-switch biases. One possible explanation for the Transition bias reset after error trials is that, when the prediction about the recent history of rewarded repetitions and alternations fails, animals downplay the weight of the prior and prioritize the sensory information by paying more attention to the sound after error choices. Our current set of experiments in the Silent task suggest otherwise, as the behavior of the animals still shows this reset after error even in the absence of guiding sensory stimuli. Recent modeling work using recurrent neural networks has proposed that the after-error reset could be a general adaptive phenomenon reflecting the intrinsic information asymmetry that more natural environments may generate between rewarded and non-rewarded actions (Molano-Mazon et al., 2021). Future experiments using foraging tasks with other types of spatio-temporal correlations across trials will help establish the generality of this after-error reset.

## METHODS

### Animal Subjects

Animals were male Long-Evans rats (Charles River), pair-housed during behavioral training and kept in stable conditions of temperature (23 °C) and humidity (60%) with a constant light-dark cycle (12h:12h, experiments conducted during light phase). Rats had *ad libitum* food, but water was restricted during behavioral sessions. Animals had *ad libitum* water on days with no experimental sessions. All experimental procedures were approved by the ethics committee (Comité d’Experimentació Animal, Universitat de Barcelona, Spain, Ref 390/14).

### Behavioral tasks

#### Auditory task

Rats in Group #1 (n = 29) performed an intensity level difference (ILD) categorization 2AFC task. Briefly, at each trial, an LED on the center port indicated that the rat could start the trial by poking in (Fig. 2A). After a fixation period of 300 ms, the LED switched off and two speakers positioned at both sides of the box played simultaneously an up to 1 s long AM white noise. Rats had to discriminate the side with the loudest sound (Right/Left) and seek reward in the associated port. Animals could respond any time after stimulus onset, but withdrawal from the center port *during* stimulus presentation immediately stopped the sound. Withdrawal from the center port *before* the fixation period reinitiated the trial (fixation break). After a fixation break, rats were allowed to initiate fixation again, and as many times as necessary until fixation was complete (indicated by center LED offset). Stimulus strength varied randomly from trial to trial, and the stimulus sequence was predetermined (see section ‘Stimulus sequence’) in repetitive (P_REP_ = 0.8) and alternating blocks (P_REP_ = 0.2).

#### Auditory task with Silent catch trials

Rats in Group #2 (n = 8) and Group #4 (n = 6) performed an ILD categorization 2AFC Auditory task with a random 5-10% of Silent catch trials. For Sound trials, details of the task are identical to the Auditory task. If it was a Silent catch trial, no sound was presented, but rats could still respond any time after the fixation period, as in Sound trials. Fixation breaks were treated as in the Auditory task. Stimulus sequence was also predetermined in repetitive (Group #2: P_REP_ = 0.8; Group #4: P_REP_ = 0.95) and alternating blocks (Group #2: P_REP_ = 0.2; Group #4: P_REP_ = 0.05).

#### Foraging task

Rats in Group #3 (n = 7) and Group #5 (n = 4) performed a non-perceptual 2AFC task. The beginning of the task was identical to the Auditory task (Fig. 4A). After a fixation period of 300 ms, the LED switched off and rats had to guess at which of the two lateral ports the reward would appear. Fixation break dynamics were identical to the Auditory task. Stimulus sequence was predetermined in repetitive and alternating blocks (Group #3: P_REP_ = 0.8/0.2; Group #5: P_REP_ = 0.95/0.05). Three of the rats in Group #3 were previously trained on the Auditory task (LE23-25), while the remaining four were directly trained in the Foraging task (LE56-59, and thus naïve to the Auditory task). No statistical differences were observed in any variable between animals already trained in the task and animals naïve to the task (data not shown).

#### Silent accidental trials

Rats in Group #6 (n = 18) performed an ILD categorization 2AFC task (see Auditory task), but only trials that accidentally did not produce any sound were evaluated.

In all tasks, correct responses were rewarded with a 24 µL drop of water and incorrect responses were punished with a 2 s time-out. Trials in which the rat did not make a side poke response within 8 s after leaving the center port were considered invalid and excluded from the analysis (average of 0.4% invalid trials per animal). Animals were trained one session per day lasting 50-75 min, 6 days per week, during up to 18 months. All the experiments were conducted in custom-made operant conditioning cages, the behavioral set-up controlled by an Arduino-powered device (BPod v0.5, Sanworks, LLC, Stony Brook, NY, USA) and the task was run using the open-source software PyBPod (pybpod.com). Some animals from Group #1 (LE42-47, LE54-55 and LE76-81) were evaluated for other purposes in (Hernández-Navarro et al., 2021). Animals from group #2, #3, #4 and #5 were also used in Molano-Mazón et al., 2021.

| | # rats | # sessions<br>(mean ± std) | # trials<br>(mean ± std) | # Silent trials<br>(mean ± std) |
| --- | --- | --- | --- | --- |
| Group #1 (Sound, 80/20)<br>LE36-41, LE42-47, LE54-55,<br>LE76-81, LE84-86, LE101-105 | 29 | 47 ± 23 | 27,119 ± 14,953 | --- |
| Group #2 (Catch, 80/20)<br>LE42-47, LE54-55 | 8 | 64 ± 28 | 39,299 ± 18,263 | 3,235 ± 1,431 |
| Group #3 (Foraging, 80/20)<br>LE23-25, LE56-59 | 7 | 19 ± 1.2 | 14,380 ± 3,104 | 14,380 ± 3,104 |
| Group #4 (Catch, 95/5)<br>LE76-81 | 6 | 16 ± 0.8 | 7,666 ± 755 | 762 ± 87 |
| Group #5 (Foraging, 95/5)<br>LE56-59 | 4 | 52 ± 7 | 31,190 ± 8,703 | 31,190 ± 8,703 |
| Group #6 (Accidental, 80/20)<br>LE36-41, LE42-47, LE101-105 | 18 | 60 ± 18 | 35,904 ± 12,071 | 1,725 ± 1,031 |

### Acoustic stimulus

In the intensity level discrimination 2AFC task, the stimulus *S_k_(t)* in the *k*-th trial was created by simultaneously playing two amplitude modulated (AM) sounds *T_R_(t)* and *T_L_(t)*:

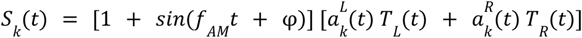

The AM frequency was *f*_AM_ = 20 Hz and the phase delay φ = 3π/2 made the envelope zero at *t* = 0. *T_L_(t)* and *T_R_(t)* were broadband noise played either from the left or the right speaker, respectively. The amplitudes of the sounds *T_L_(t)* and *T_R_(t)* were calibrated at 65 dB SPL using a free-field microphone (Med Associates Inc, ANL-940-1). Sounds were delivered through generic electromagnetic dynamic speakers (ZT-026 YuXi) located on each side of the chamber.

### Stimulus Sequence

A two-state Markov chain generated a sequence of stimulus category *c_k_* = {-1,1}, which determined whether the reward in each trial was available in the left or right port, respectively (Fig. 3A). The Markov chain was parameterized with a single transition probability, P_REP_, which quantified the probability to repeat the previous stimulus category (Fig. 3A). The probability P_REP_ varied in blocks of 80 trials between P_REP_=0.8 (Rep) and P_REP_=0.2 (Alt). In Group #4 the probabilities were made P_REP_=0.95 (Rep) and P_REP_=0.05 (Alt). The stimulus category *c_k_* set which of the sounds, *T_L_(t)* or *T_R_(t)*, was dominant, which determined the statistics of the sound amplitudes 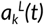 and 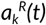. In each trial, independently of the stimulus category (*c_k_*), the stimulus strength (*s_k_*) was also randomly generated. Stimulus strength *s_k_* defined the relative weights of the dominant and non-dominant sounds: i.e. when *s_k_* = 1, only the dominant sound was played (easy trial), whereas when *s_k_* = 0 the two sounds had on average the same amplitude (hard trial). We used four possible values for s = 0, 0.25, 0.5 and 1. The stimulus evidence was defined in each trial as the combination *e_k_* = *c_k_*\**s_k_*, thus generating seven different stimulus evidence values *e_k_*=(0, ±0.25, ±0.5, ±1). The value of *e_k_* determined the p.d.f. from which the two sets of instantaneous evidences 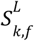 and 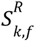 were drawn at each 50 ms frame *f* (Hermoso-Mendizabal et al., 2020). When *e_k_*: = ±1 the p.d.f. for 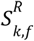 was *f*(*x*)=δ(*x*∓1) (i.e., a Dirac delta p.d.f.), whereas when *e_k_* ∈ (−1,1), it was a stretched beta distribution with support [−1,1], mean equal to *e_k_* and variance equal to 0.06. The p.d.f. Distribution for 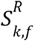 was the mirror image with respect to zero (i.e. a Dirac or beta distribution with mean −*e_k_*). Finally, the amplitudes 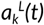 and 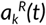 of the two AM envelopes were obtained using:

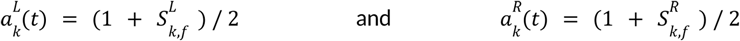

with *f* referring to the frame index that corresponds to the time *t*.

### Psychometric curve analysis

In the Auditory task, we computed the repeating psychometric curve for each animal, by pooling together trials across all sessions separately for each type of block (Rep or Alt) and using all the seven stimulus evidence values e = 0, ±0.25, ±0.5, ±1. We calculated the proportion of repeated responses as a function of the repeating stimulus evidence (*ê*) defined for the *t*-th trial as *ê_t_* = *r_t_*_-1_*e_t_*, with *r*_t−1_ = {−1,1} representing the previous response (i.e. left or right, respectively). Thus, positive values of *ê* denote trials in which animals had evidence to repeat their previous choice, while negative values of *ê* in which they had evidence to alternate. Psychometric curves were fitted to a 2-parameter probit function (using Matlab function nlinfit):

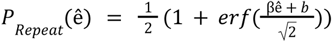

The sensitivity β quantified the stimulus discrimination ability, while the parameter *b* was defined as the *repeating bias* which captured the animal preference to repeat (*b* > 0) or alternate (*b* < 0) their previous reponse. Across-subject error bars, corresponded to the 1^st^ and 3^rd^ quartiles.

To quantify the tendency to repeat or alternate in trials with no stimulus (i.e. Silent catch trials or trials in the Foraging task), as no psychometric curve could be drawn, we used the Repeating Probability simply defined as the probability to repeat the previous choice. We also calculated separately for each block type the Excess of Repeating Probability as *ERP* = ±*(P_rep_* − 0.5) with the positive sign used for the Repeating block and the negative sign for the Alternating block. Hence for example an ERP = 0.2 meant an excess of 0.2 probability points to repeat in the Repeating block and an excess of 0.2 probability points to alternate in the Alternating block.

### Generalized linear model (GLM) analysis

#### Net evidence GLM

We fitted a GLM to quantify the weight of different features - such as the current stimulus and previous history events-had on the choices of the animals (Busse et al. 2011; Frund, Wichmann, and Macke 2014; Abrahamyan et al. 2016; Braun, Urai, and Donner 2018; Hermoso-Mendizabal et al. 2020). The probability that the response *r_t_* in trial *t* was to the right was modeled as a linear combination of the features passed through a probit function:

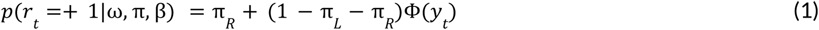

In which π_R_ and π_L_ represent the lapse rates for left and right responses, the probit function Φ(*x*) is the cumulative of the standard normal function and its argument in trial *t* reads:

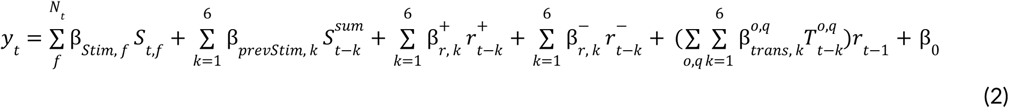

The current stimulus, given by 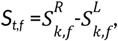 represents the intensity difference between the Right and the Left sounds in each frame *f*, with *f* = 1,2…N_t_ and *N_t_* being the number of frames listened in trial *t*. The contribution of trial history included the impact of the previous ten trials (t-1, t-2, t-3…; grouping the impact of trials t−6 to trial t−10 in one term). The previous stimuli, given by 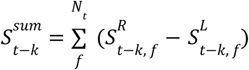, representing the intensity difference between the Right and the Left sounds summed across the *N_t_* listened frames. The terms 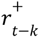 represented the previous correct responses being −1 (correct left), +1

(correct right), or 0 (error response). Similarly, 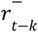 represented previous incorrect responses. Previous transitions 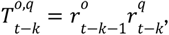, which were separated into four different types {*o, q*} = {+, +}, {+, −}, {−, +}, and {−, −}, depending on the outcomes of trial *t−k* (q) and *t−k−1* (o) (Hermoso-Mendizabal et al., 2020).

#### Interaction GLM

To evaluate if the presentation of previous stimuli impacts the perception of the current stimulus, we added an additional interaction term defined as the product of the current and previous stimuli. To do this, we introduced three variations with respect to the Net evidence GLM. First, we separated the contributions of the current and previous stimuli, previously defined as the net evidence, into left 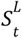 and right 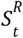 contributions. Second, in order to reduce the amount of regressors, we grouped the Current stimulus regressors, previously defined for each sample (Eq. 2), into a single regressor defined as an exponentially weighted sum of each of the listened frames: 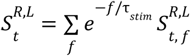. The previous stimuli were also grouped into a single regressor that captured the weighted sum of all previous left or right stimuli: 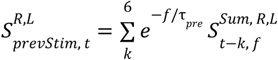. The decay time constants τ_stim_ and τ_prev_ were calculated by fitting an exponential to the individual weights *β_Stim, f_* (across frames) and *β_prevStim, k_* (across trial lag) obtained from the *Net evidence GLM*, respectively. The averages of the individual constants were τ_stim_=1,54 frames and τ_prev_ = 3,88 trials. Third, we included ipsilateral interaction terms relating previous and current choices: 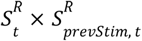 and 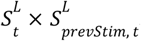. Contralateral interactions were not included for simplicity. We these modifications, the argument of the probit function now reads:

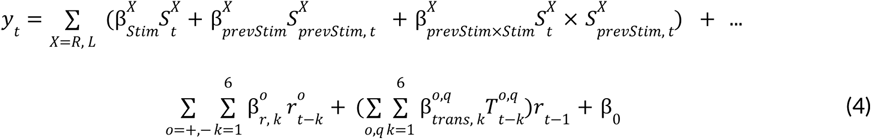

### Repeating bias recovery

We calculated the baseline repeating bias *b* of each block (Rep or Alt) in both tasks (Sound and Foraging) by averaging the repeating probability during the last 5 trials of each block. Then, we compared the repeating bias trial by trial for the first 15 trials of a block to check how many trials it takes to return to baseline levels.

### Reaction times

Reaction time (RT) was defined as the period between stimulus onset and the center port withdrawal (Fig. 2A and 4A). Fixation breaks (FB) were defined as withdrawals from the center port during the fixation period (300 ms). Only the first center port withdrawals of each trial, either FB or RT, were analyzed: after a FB, further FBs and the subsequent valid response were discarded to remove possible serial effects within a single trial (mean ± standard deviation, 16±5 % of total withdrawals). RTs longer than 1s after fixation onset were removed from the analysis (0.5±0.6 % of total withdrawals).

## Data code availability

Data and code will be made available on a public repository once the manuscript is published.

## Acknowledgements

We thank Daniel Linares for comments on the manuscript, Manuel Molano, Lluís Hernández-Navarro and Alex Hyafil for useful discussion, and Lejla Bektic and Lorena Jiménez for help with training of the animals. This research was supported by the Spanish Ministry of Economy and Competitiveness together with the European Regional Development Fund (IJCI-2016-29358 to D.D.; RTI2018-099750-B-I00 to J.R.) and the European Research Council under the European Union’s Horizon 2020 research and innovation program (ERC-2015-CoG - 683209 PRIORS to J.R.). Part of this work was developed at the building Centre Esther Koplowitz, Barcelona.

## Author contributions

D.D. and J.R. conceived and designed the project and experiments; D.D. carried the experiments and analyzed the experimental data; D.D., and J.R. interpreted the data and wrote the manuscript.

## SUPPLEMENTARY FIGURES

**Figure 1 - Fig. supplement 1.**
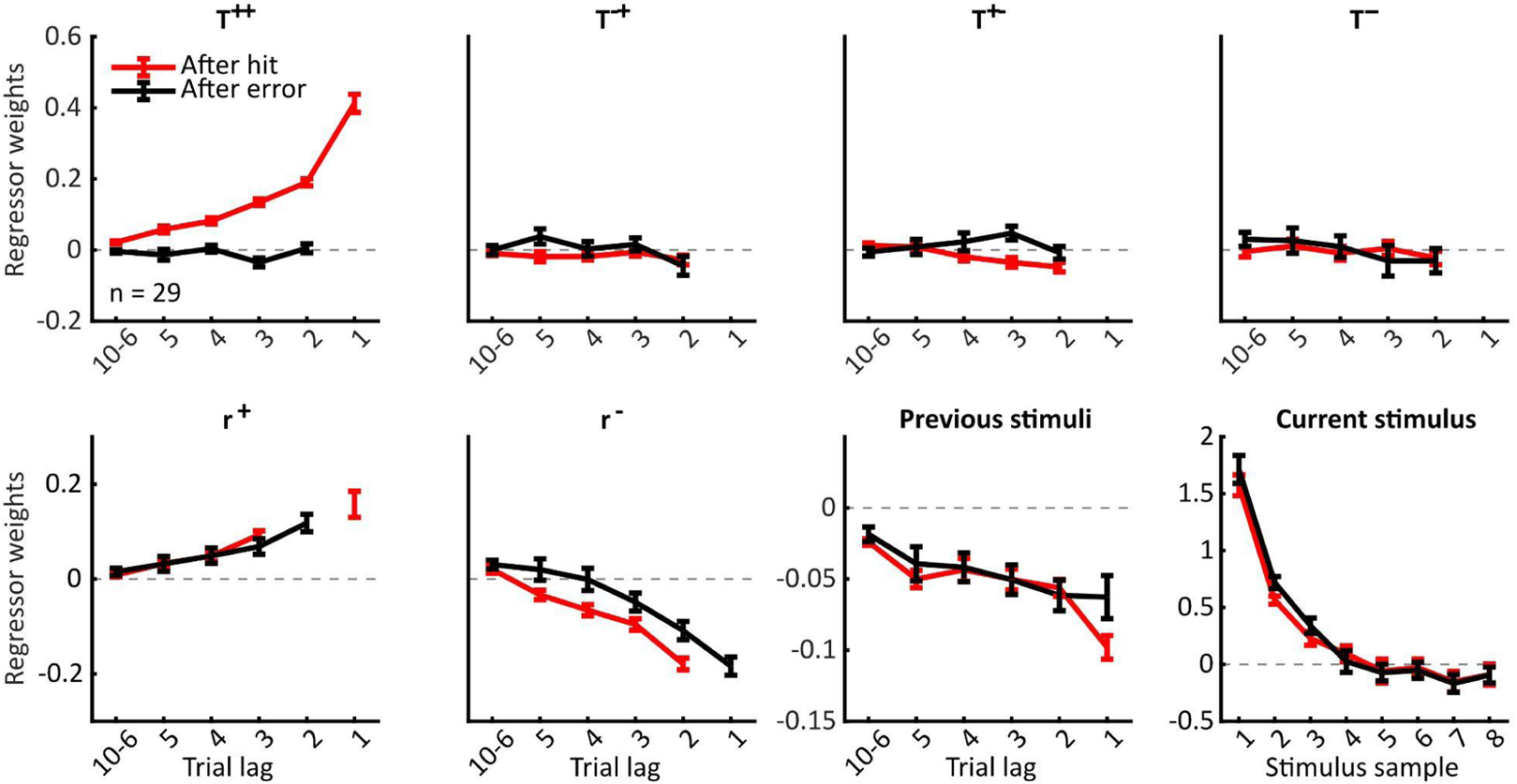
**Full GLM in the Auditory task.** We fitted the data separately for after-correct (red) and after-error trials (black). We assessed the contribution of the predictive transition bias (T^++^, two consecutive correct transitions; T^−+^, a sequence of an incorrect and a correct transition; T^+−^, a sequence of an correct and an incorrect transition; T^−−^, two consecutive incorrect transitions) and correct rewarded and unrewarded responses, respectively (r^+^, reflecting *win-stay* behavior; r^−^, reflecting *lose-switch* behavior). A correct transition may be either a correct repetition or alternation. Both biases are then combined with the stimulus evidence of previous trials (Previous stimuli) and of the current trial (Current stimulus) to generate a choice. Labels described as in Figure 1. For simplicity, and because the impact of transitions with an incorrect choice is barely noticeable, T^+−^, T^−+^, and T^−−^ will not be analyzed. After-correct choices, the decisions of the animals were positively and strongly modulated by the Current stimulus and also positively, although less strongly, by the previous correct transitions (T^++^). Previous correct (r^+^) and incorrect responses (r^−^) also modulate the rats’ decisions, but to a lesser extent than the predictive transition bias. Previous stimuli had a small but consistent negative impact on choices. The weight of previous correct transitions (T^++^) reset after-error.

**Figure 1 - Fig. supplement 2.**
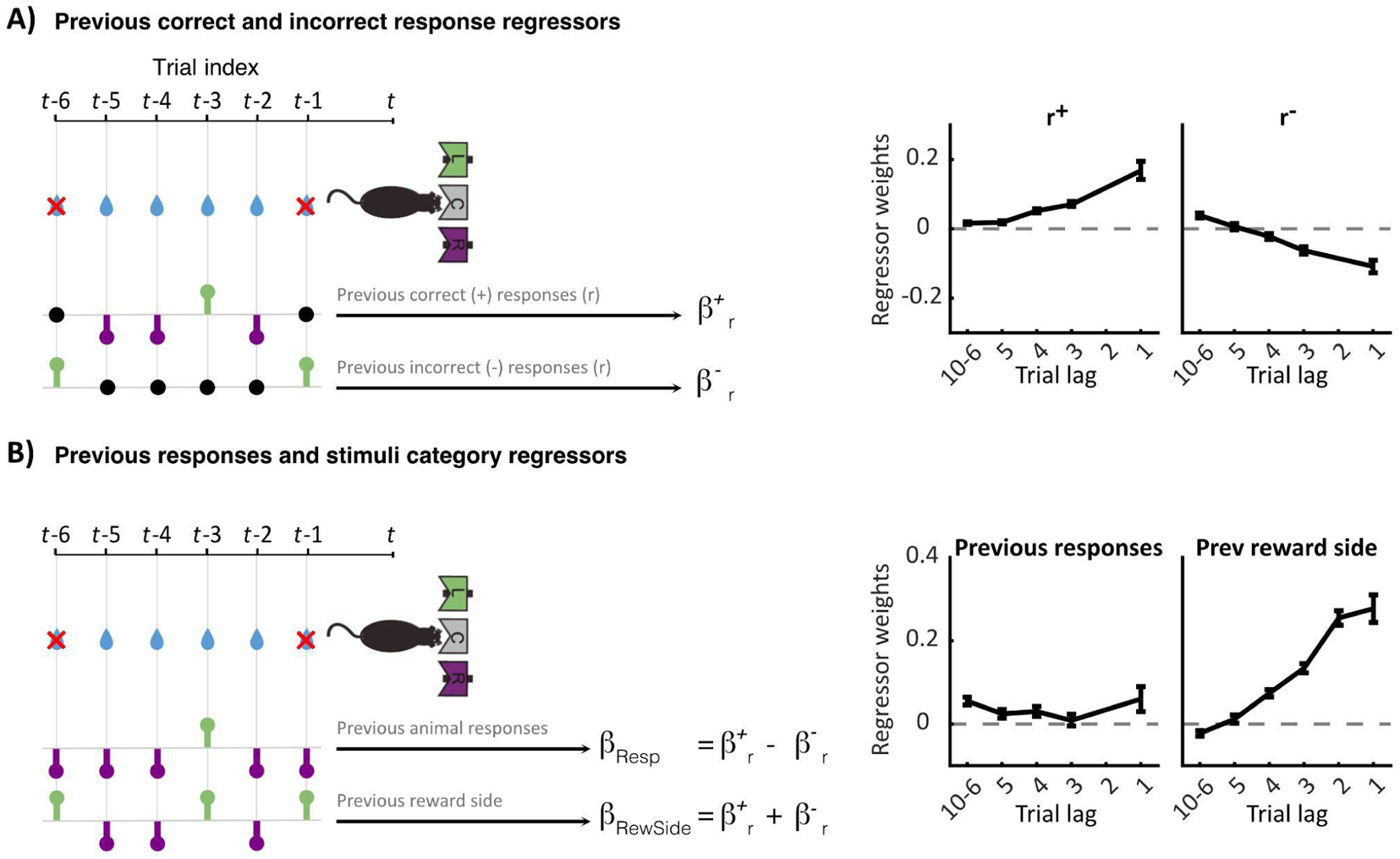
**Past choices and outcomes regressors.** There are two ways of representing the past choices and their outcomes, as **(A)** correct rewarded and unrewarded responses (r^+^ and r^−^) or as **(B)** previous responses and previous reward side (Resp, RewSide). **A)** Previous correct responses are labeled −1 for after correct Left, 1 for after correct Right and 0 for after errors. Previous incorrect responses are labeled as 0 after correct, −1 for after incorrect Left and 1 for after incorrect Right. **B)** However, previous responses r^+^ and r^−^ can be re-parameterized as Previous responses after a subtraction (r^+^ − r^−^) and as Previous reward side after a sum (r^+^ + r^−^). In that case, both regressors may take just two values ([−1,1]). Such re-parametrization shows that, although correlated, Previous reward side and Previous stimuli are separable in the model because while Previous reward side is binary ([−1 for left, 1 for right]), Previous stimuli are graded.

**Figure 1 - Fig. supplement 3.**
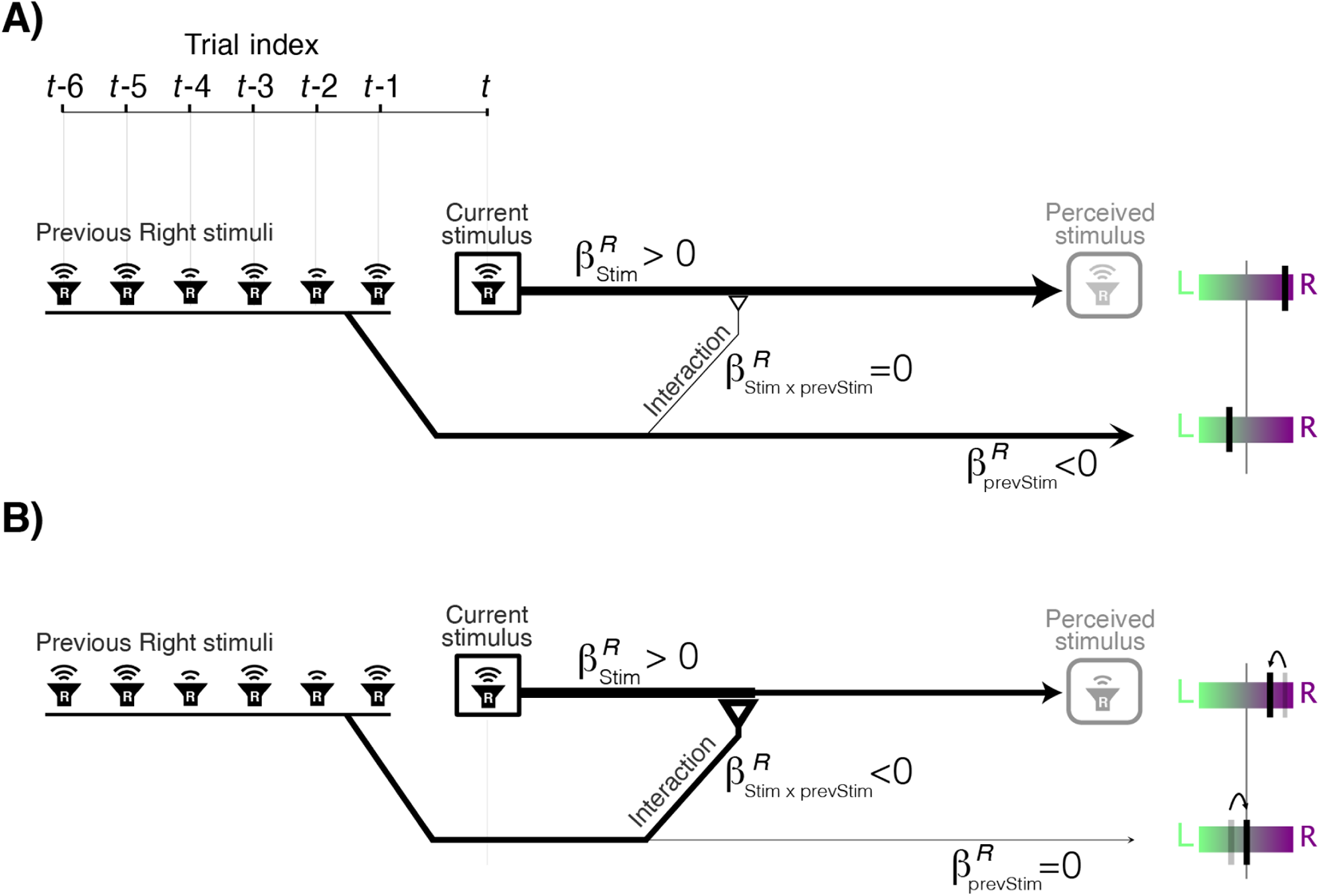
Alternative mechanisms underlying the Previous Stimulus repulsive bias and their characterization using the Interaction GLM. A) Schematic showing the case in which the Previous Stimulus repulsive bias does not affect the perception of the Current stimulus. In this scenario, the weight of the direct impact of Previous stimuli coming from the Right is negative (β^R^_prevStim_<0) and has the opposite sign of the weight of the Current Stimulus coming from the Right (β^R^_Stim_>0). The stimuli coming from the Left (not shown) exhibit the same opposing behavior but with the reverse sign as they decrease the probability to choose Right, i.e. β^L^_Stim_<0 and β^L^_prevStim_>0. Because previous stimuli do not modulate the impact of the current stimulus, the interaction weight is null (i.e. β^R^_Stim × prevStim_=0; same for the Left side). Color bars on the right illustrate the direct impact of Previous (top) and Current (bottom) stimuli shown in the example. B) Case in which the Repulsive bias arises from the modified perception of the Current stimulus. In this scenario, Previous stimuli have no *direct* impact on choice (i.e. β^R^_prevStim_=0) but cause repulsion by a negative interaction weight (β^R^_Stim × prevStim_<0) which decreases the final weight of the current Stimulus (notice how the width of the Current Stimulus arrow decreases after the interaction yielding a reduced “*perceived stimulus*”) .

**Figure 2 - Fig. supplement 1.**
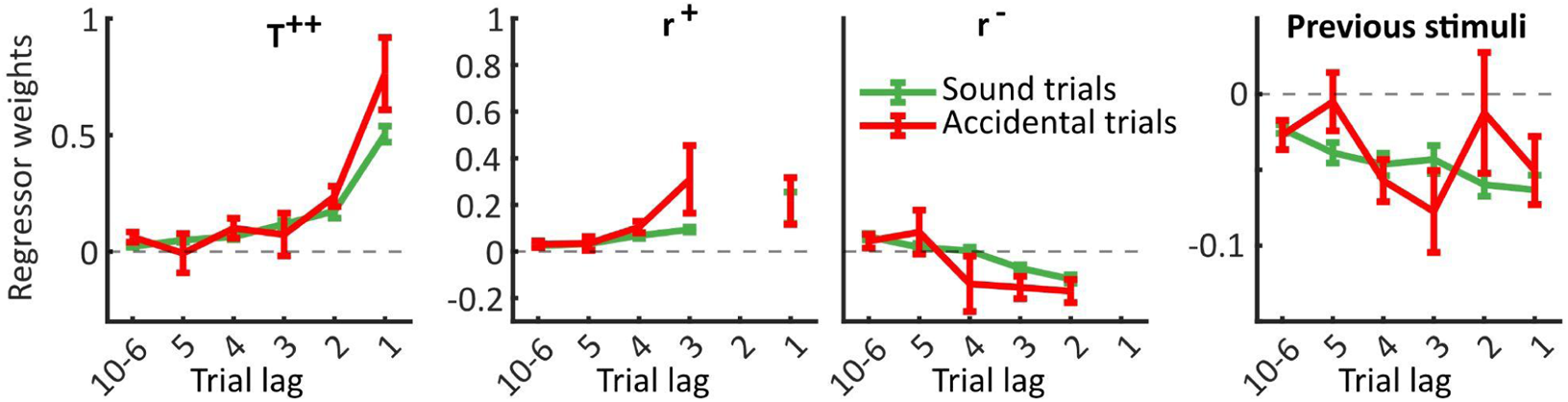
**GLM for Accidental silent trials.** We fitted the data separately for Sound (green) and Accidental silent trials (red) for after-correct trials. We confirmed that the Aftereffect bias was also present in Accidental silent trials, i.e., trials in which our set-up failed to produce a sound (Previous stimuli; ‘trial’ × ‘lag’ interaction: F_1,195_ = 0.9, p = .3427; ‘trial’ main effect: F_1,195_ = 0.94, p = .3335). The significant larger weights for T^++^ and the decrease for both r^+^ and r^−^ can be explained by the fact that Accidental silent trials normally occurred for trials with short reaction times, which coincide with trials with high repeating bias *b* - thus higher weights for the T^++^ and smaller for r^−^ regressors.

**Figure 3 - Fig. supplement 1.**
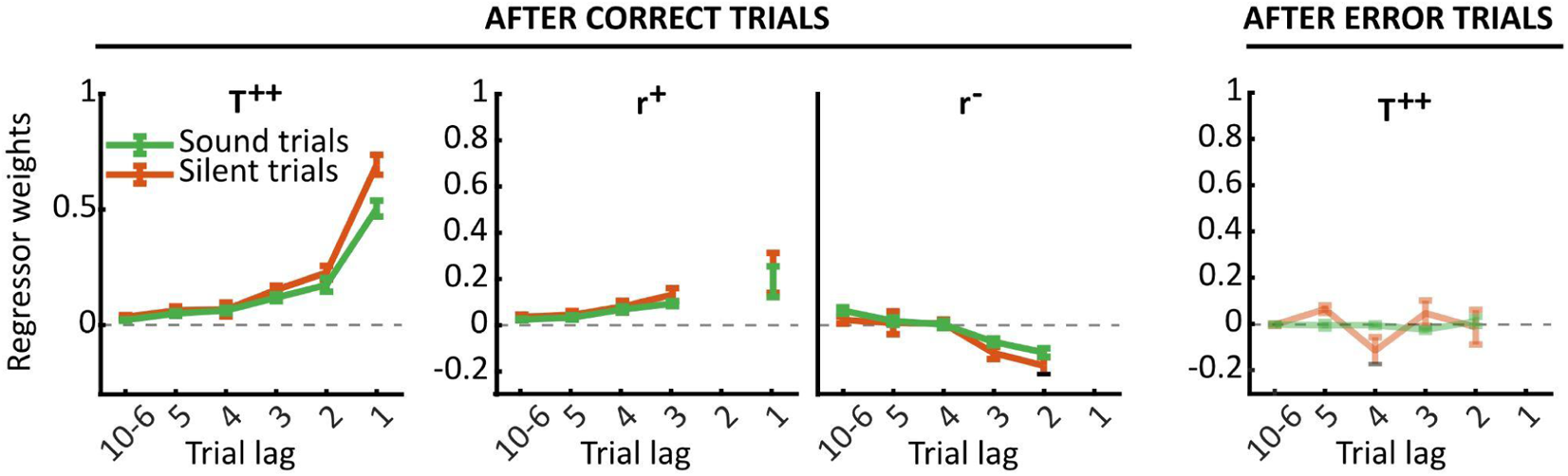
Comparison of the effect of previous correct and incorrect responses, and the reset after-error in Sound and Silent catch trials. We fitted the data separately for Sound (green) and Silent trials (orange) for after-correct (three left charts) and after-error trials (right chart). We assessed the contribution of the predictive transition bias (T^++^) and previous correct and incorrect responses (r^+^; r^−^), as described in Figure 1 - Fig. supplement 1. After-correct choices, no differences were observed between Sound and Silent trials neither for previous correct (r^+^; ‘trial’ × ‘lag’ interaction: F_1,69_ = 0.23, p = .632; ‘trial’ main effect: F_1,69_ = 0.71, p = .4033) nor incorrect responses (r^−^; ‘trial’ × ‘lag’ interaction: F_1,69_ = 0.46, p = .4997; ‘trial’ main effect: F_1,69_ = 1.68, p = .1988). The weight of previous correct transitions (T^++^) reset after-error in both Sound and Silent trials, and the weights are not different between them (T^++^ after-error; ‘trial’ × ‘lag’ : F_1,69_ = 0.07, p = .7896; ‘trial’: F_1,69_ = 0.06, p = .8124).

**Figure 3 - Fig. supplement 2.**
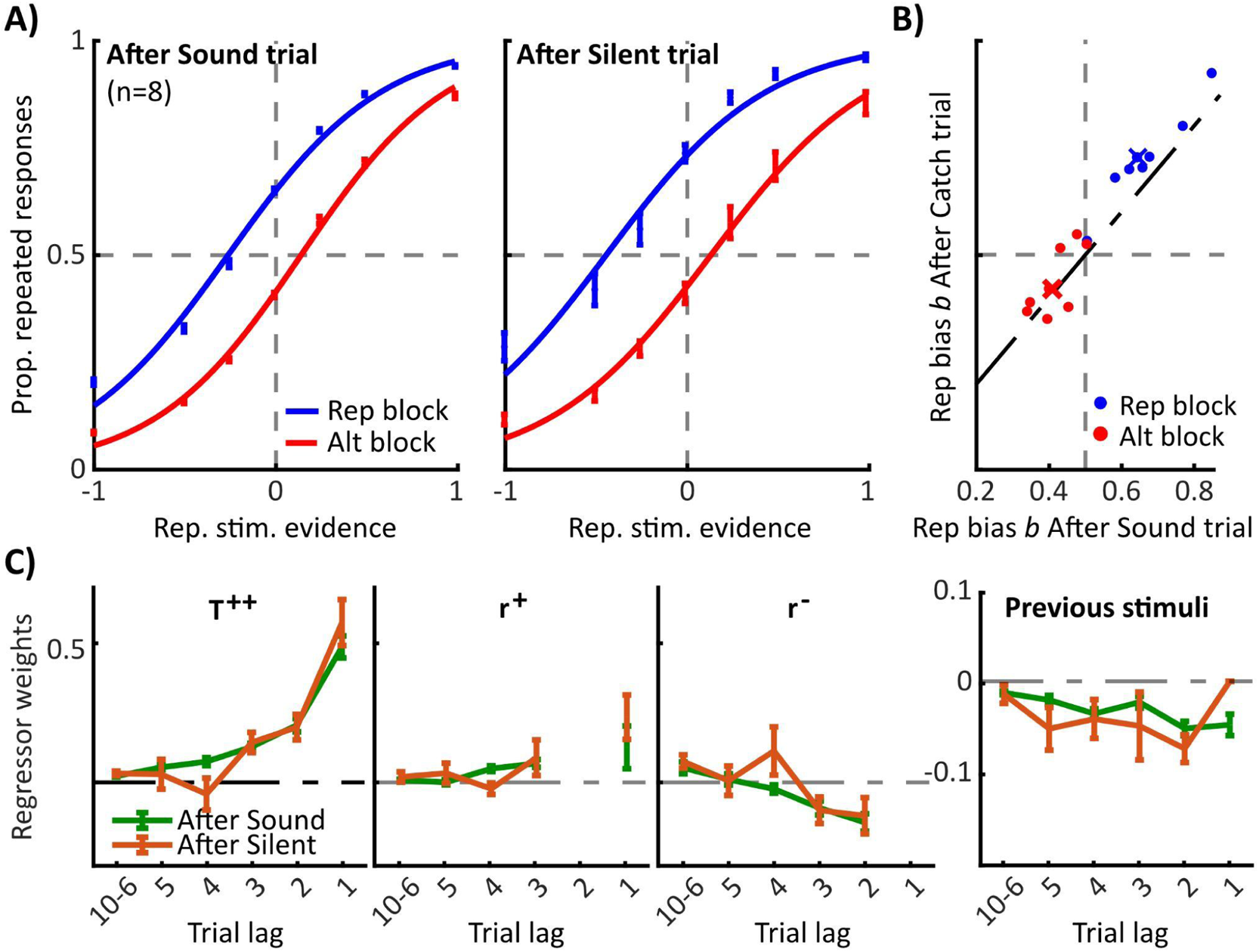
History biases do not reset when the previous trial is silent. **A)** Mean psychometric curves (n = 8 rats) showing the proportion of repeated responses computed in trials following a correct Sound trial (left) or a correct Silent trial (right), sorted by blocks (blue: repetitive; red: alternating). **B)** Repeating bias *b* After Silent trials in Rep blocks is enhanced as compared to After Sound trials (0.74±0.09 vs 0.68±0.09; paired t-test, p < .0001), while there is no difference in Alt blocks (0.46±0.07 vs 0.43±0.05; p = .0609). **C)** Influence of past events on current choice After Sound (green) or After Silent trials (yellow). Same convention as in Figure 3. Notice that weights for the Previous stimuli kernel go to zero at lag −1 in After Silent trials (orange line). No differences between After Sound and After Silent trials were observed neither in transition bias (T^++^: ‘trial’ × ‘lag’ interaction: F_1,85_ = 0.07, p = .7888; main factor ‘trial’ F_1,85_ = 0.09, p = .7674) nor in r^−^ (‘trial’ × ‘lag’ interaction: F_1,69_ = 0.31, p = .5812; ‘trial’ F_1,69_ = 0.83, p = .3647). A small significant difference was found between After Sound and After Silent trials in r^+^ (‘trial’ × ‘lag’ interaction: F_1,69_ = 3.99, p=0.0498; ‘trial’ F_1,69_ = 3.67, p=0.0596), suggesting animals have a larger trend to repeat previous successful decisions after the sudden absence of sensory inputs.

**Figure 5 - Fig. supplement 1.**
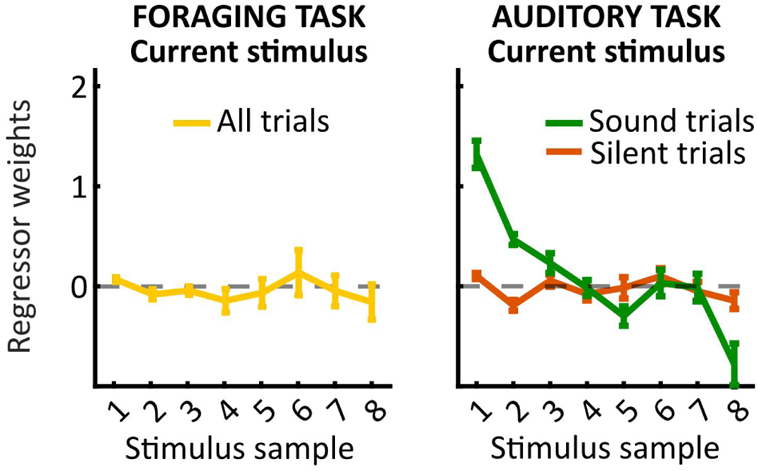
**Kernels for Current stimulus in both Foraging and Auditory tasks.** The sensory information in a Sound trial in the Auditory task (green line) can last between a few ms and up to 1 s, the maximum duration of a played sound. For simplicity, and as most of rats’ reaction times are shorter than 200 ms, we only plot 8 sound samples of 50 ms each. For both the Foraging task trials (yellow line) and the Silent trials in the Auditory task (orange line), the sound is not played and the kernels for the current trial are flat. Silent trials in the Auditory task allow us for a direct comparison of the history choice biases in the two tasks, as the Current stimulus kernel in both tasks is zero.

**Figure 5 - Fig. supplement 2.**
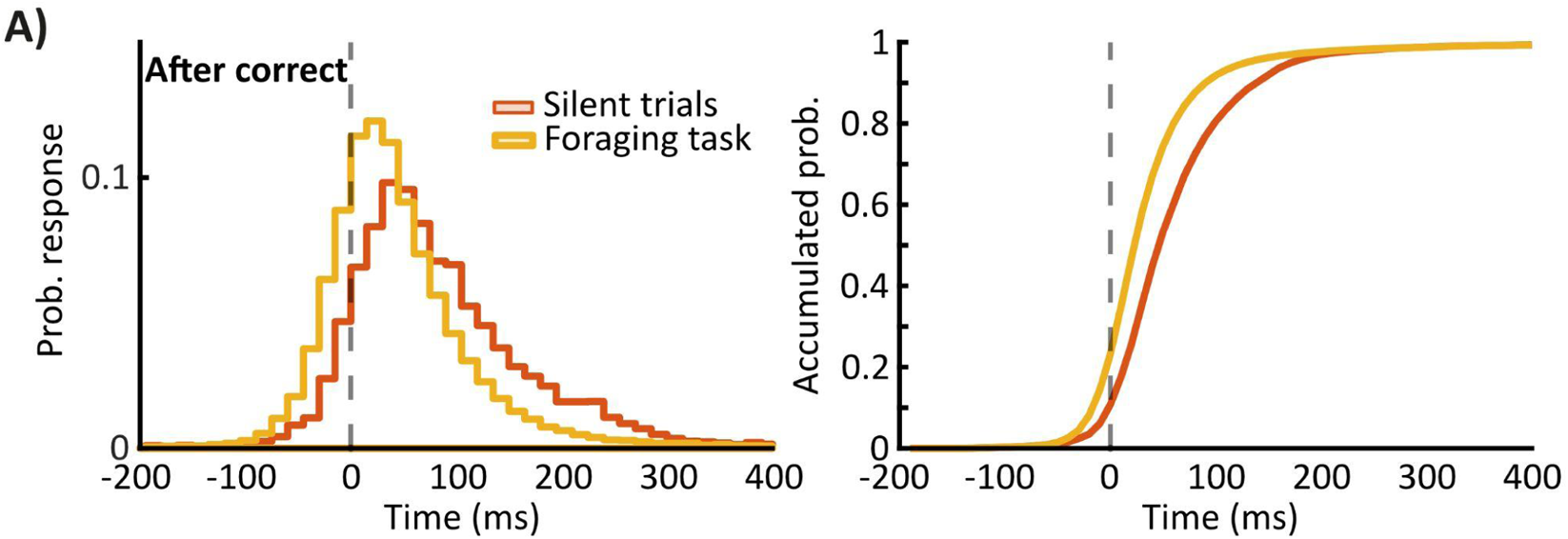
Reaction times differences between the Sound and the Foraging task. **A)** Left panel, RT distributions for Silent trials in the Auditory task (orange; n = 8 rats) and for the Foraging task (yellow, n = 7 rats). **B)** RT cumulative distribution function (pdf) for the trials in the Foraging task and for the Silent trials in the Auditory task. RTs for the Foraging task (yellow line: mean ± std 53 ± 110 ms) were shorter than RTs for Silent trials in the Auditory task (orange line: 92 ± 131 ms; Kolmogorov-Smironov test, KS score = 0.23, p < .0001). Rats in the Foraging task only prioritized responding as fast as possible. The immediate consequence of such a decision is an increase in the number of fixation breaks (which reinitiate the trial) with 25.91% of the 77,543 trials in the Foraging task as compared to 10.77% of the 27,236 Silent trials in the Auditory task. The proportion of fixation breaks was significantly different between Silent catch trials in the Auditory task and the Foraging task trials (X^2^, 1 d.f. = 2691.47, p < .0001).

**Figure 5 - Fig. supplement 3.**
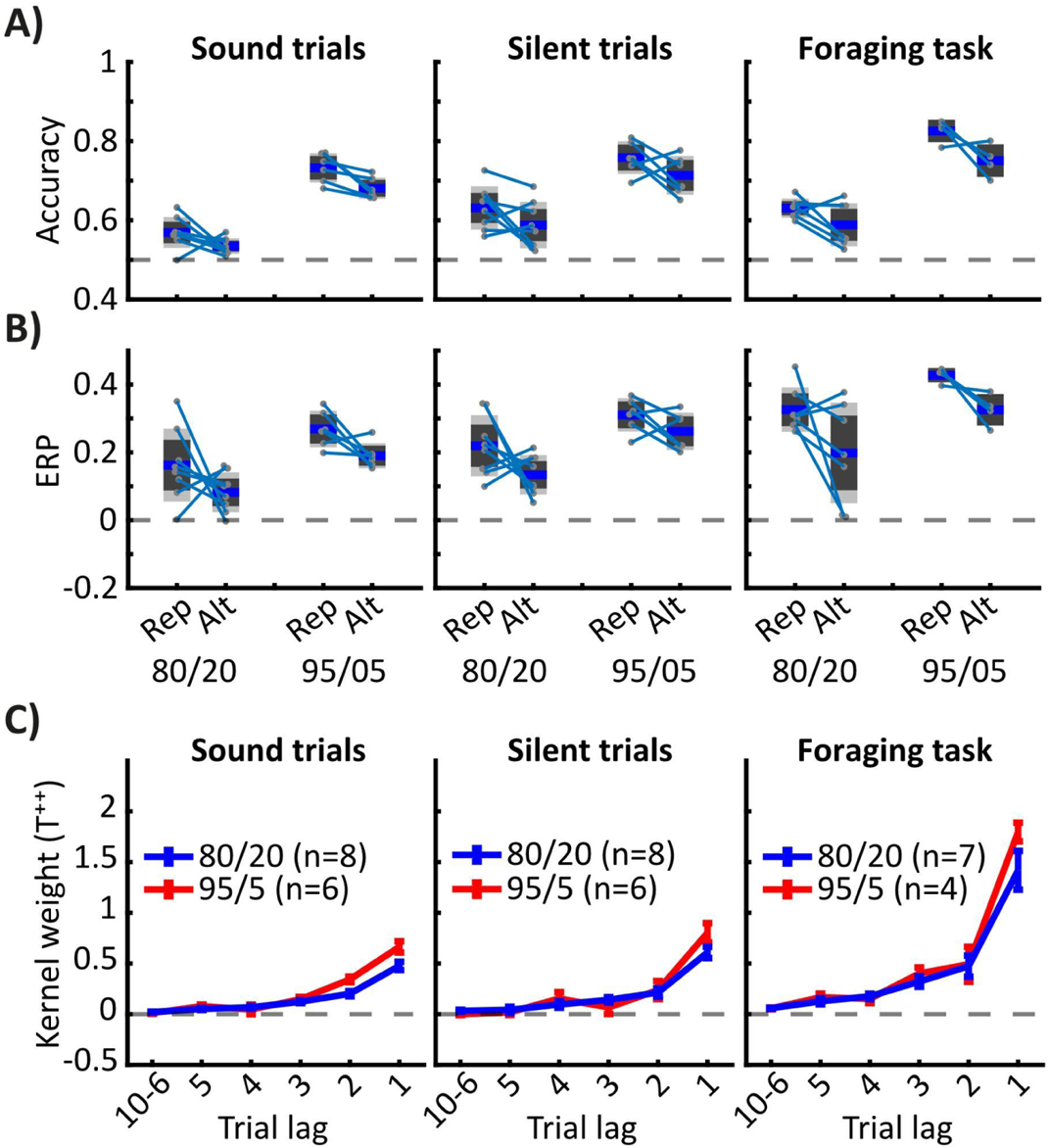
**Effect of the predictability of the sequential correlations on history biases.** We trained a new batch of animals (Group #3, n = 6 rats) in the same Auditory task shown in Fig. 2 and we retrained 4 animals from Group #2 in the same Foraging task shown in Fig. 4, but with more estable sequential correlations (Rep, P_REP_ = 0.95; Alt, P_REP_ = 0.05). **A)** Accuracy for Sounds trials and Silent trials in the Auditory task, and all trials in the Foraging task sorted by block (Rep/Alt) and environment (Regular: P_REP_=0.8/0.2; Extreme: P_REP_=0.95/0.05). Accuracy in the extreme environment was larger than in the regular environment (3-way ANOVA for ‘accuracy’ with ‘trial’ (Sound/Catch/Silent), ‘block’ (Rep/Alt) and ‘environment’ (8020/9505) as variables; ‘environment’ F_1,66_ = 240.1, p < .0001), independently of other factors (triple interaction F_2,66_ = 0.2, p = .8204; ‘environment’ × ‘trial’ F_2,66_ = 2.25, p = .1133; ‘environment’ × ‘block’ F_2,66_ = 0.76, p = .3856). **B)** Exces of repeating probability (ERP) is larger for the extreme environment than for the regular environment (‘environment’ F_1,66_ = 37.01, p < .0001), and such difference does not depend on other variables (triple interaction F_2,66_ = 0.09, p = .9109; ‘environment’ × ‘trial’ F_2,66_ = 0.01, p = .9873; ‘environment’ × ‘block’ F_1,66_ = 0.34, p = .5596). **C)** Comparison of the T^++^ kernels obtained in the regular (P_REP_=0.8/0.2) and the extreme environment (P_REP_=0.95/0.05) for Sound trials (left), Silent trials (center) and the Foraging task (right). Transition bias is not enhanced in the extreme environment as compared to the regular environment (‘environment’ F_1,222_ = 2.13, p = .1463; ‘environment’ × ‘trial’ interaction F_2,222_ = 1.39, p = .2522; triple interaction F_2,222_ = 0.73, p = .4817).

